# Mucosal ACP subunit vaccine formulated with a small molecule STING agonist drives protective T_H_1/T_H_17 responses and serum bactericidal antibodies against *Neisseria gonorrhoeae*

**DOI:** 10.64898/2026.11.02.763630

**Authors:** Abhishek Chanda, Ines A. Cadena, Junaid Nazir, Laura Springgay, Ian P. Rose, Jinu Abraham, Chenwei Lin, Mansab Ali Saleemi, Andrew N. Macintyre, Victor DeFilippis, Aleksandra E. Sikora

## Abstract

STING agonists are emerging vaccine adjuvants; however, only cyclic dinucleotides (low permeability and stability) have been tested against pathogenic bacteria. We examined diABZI, a non-nucleotide STING agonist active in human and murine orthologs, combined with the *Neisseria gonorrhoeae* (Ng) Adhesin Complex Protein, a conserved adhesin and human lysozyme C inhibitor, as a gonococcal vaccine. Intranasal administration of recombinant ACP (rACP)+diABZI accelerated gonococcal clearance, in contrast to rACP or rACP+CpG+MPLA, despite comparable antibody responses. While both diABZI and CpG+MPLA induced T_H_1, diABZI also induced T_H_17 polarization. Peptide mapping localized the immunodominant T cell epitopes to the ACP–lysozyme C interface. rACP+diABZI elicited higher human serum bactericidal activity (hSBA), that correlated with Ng clearance, and antibodies that reversed human lysozyme C inhibition and interfered with Ng adhesion and invasion in 3D human vaginal/urethral mucosal models. Passive serum transfer did not recapitulate protection, implicating mucosal antibodies and cellular immunity over circulating IgG.

## INTRODUCTION

Gonorrhea, caused by the Gram-negative bacterium *Neisseria gonorrhoeae* (Ng), is the second most prevalent sexually transmitted infection worldwide, with an estimated 82.4 million new cases in 2020 ^1,2^. Women experience disproportionate health consequences from infection including pelvic inflammatory disease, ectopic pregnancy, and miscarriage. In men, Ng typically presents as urethritis but can ascend to the epididymis or testes ^3,4,5^. Infertility may occur in both sexes, and disseminated gonococcal infection, though uncommon, can cause systemic complications while neonatal conjunctivitis following vertical transmission can lead to corneal scarring and blindness ^4^. Amplifying its public health impact, gonorrhea increases both acquisition and transmission of HIV ^4,6^. Antimicrobial resistance continues to erode treatment options; although two new antibiotics (zoliflodacin and gepotidacin) were recently approved by the U.S. FDA for uncomplicated urogenital gonorrhea, Ng has developed resistance to every antimicrobial class introduced ^1,2,7^. No licensed vaccine exists and their development faces formidable biological barriers. For instance, Ng undergoes antigenic and phase variation and correlates of protection remain undefined. Additionally, the bacterium subverts adaptive immunity by promoting TGF-β-dependent T_H_17 polarization and IL-10-producing type 1 regulatory T cells while suppressing T_H_1 and T_H_2 effector responses that preclude establishment of protective immune memory ^8–11,12^.

The meningococcal serogroup B vaccine 4CMenB (Bexsero), a four-component formulation comprising three recombinant proteins (fHbp, NadA, NHBA) and outer membrane vesicles (OMVs) from a New Zealand meningococcal outbreak strain, showed ∼30–40% pooled cross-protection against gonorrhea in observational studies and was adopted in 2025 by the United Kingdom for targeted vaccination of gay and bisexual men ^13,14^. However, GoGoVax, the largest randomized controlled trial evaluating 4CMenB for gonorrhea prevention, found no reduction in incidence among high-risk men (vaccine efficacy –0.5% ^15,16,17^) consistent with an earlier open-label trial ^18^. These results call into question the premise of OMV-mediated cross-protection. Earlier gonococcal vaccine candidates, including killed whole-cell and pilin-based formulations, also failed to confer durable protection in clinical trials ^19^, and the next generation candidate Generalized Modules for Membrane Antigens (GMMA)-based formulation, engineered OMVs (NgG, GSK), was withdrawn from the developer’s pipeline before efficacy results were reported^1,20^. These setbacks underscore the clinical need for novel vaccine approaches.

<u>ST</u>imulator of <u>IN</u>terferon <u>G</u>enes (STING) is an adaptor molecule essential to the innate immune response to cytosolic DNA. It is endogenously stimulated by the secondary messenger 2’3’ cyclic GMP-AMP (cGAMP), synthesized by the DNA receptor cyclic GMP-AMP synthase (cGAS). Activated STING induces expression of type I interferon (IFN-I) and proinflammatory genes that generate a tissue environment enhanced for antigen processing and presentation. Importantly, pharmacologic activation of STING has been shown to augment efficacy of vaccines against viral, protozoal, and bacterial pathogens^21–24^. In a *Mycobacterium tuberculosis* murine infection model, intranasal (IN) delivery of a cyclic dinucleotide (CDN) STING agonist induced T_H_1 and T_H_17 responses, with T_H_17 cytokine production correlating with vaccine-mediated protection that exceeded that of BCG^22,23^. Similarly, a CDN-adjuvanted *Chlamydia trachomatis* antigen induced IL-17A- and IFNγ-producing CD4+ T cells and reduced *C. muridarum* shedding after vaginal challenge^24^. Importantly, this co-polarization is route-dependent: mucosal delivery drives T_H_1/T_H_17 responses, whereas parenteral administration produces T_H_1-biased or T_H_1/T_H_2-balanced immunity^23^.

No STING agonist has been evaluated in vaccines against any *Neisseria* species, and those tested against other pathogenic bacteria have been exclusively CDN-based, limited by poor membrane permeability and rapid enzymatic degradation ^25^. diABZI is a potent, non-nucleotide small-molecule STING agonist created by linking two amidobenzimidazoles to generate a synergistic compound that binds both murine and human STING and overcomes the pharmacological limitations of CDNs ^26^. diABZI has demonstrated antiviral activity against respiratory viruses and has been successfully used as a vaccine adjuvant for SARS-CoV-2 ^25^ but has not been explored for bacterial pathogens. Here, we evaluated diABZI as a mucosal adjuvant for the Ng Adhesin Complex Protein (ACP), a conserved, dual-function outer membrane protein of pathogenic *Neisseria* that acts as both an adhesin and an inhibitor of human lysozyme C^27–31^. Between 80–90% of Ng strains share a single ACP nucleotide sequence, with only 14 distinct amino acid variants identified globally, and notably, the three-dimensional structure of Ng-ACP does not incorporate any polymorphisms present in ≥1% of the global Ng population ^32^. Immunization with ACP formulated with different adjuvants generates antibodies that block ACP-mediated lysozyme inhibition ^27,33^.

Here, we report the first immunization–challenge evaluation of ACP as a gonococcal vaccine antigen and the first application of a STING agonist against any *Neisseria* species. Intranasal immunization of C57BL/6 mice with rACP adjuvanted with diABZI accelerates gonococcal clearance across three independent experiments, whereas rACP alone and rACP with the TLR9 (Toll Like Receptor 9) and TLR4 agonists CpG+MPLA fail to confer protection despite eliciting comparable systemic and mucosal antibody responses. The mechanistic basis of this divergence is selective T_H_1/ T_H_17 co-polarization: both rACP+diABZI and rACP+CpG+MPLA induce equivalent antigen-specific IFNγ (T_H_1) responses, yet only diABZI co-induces IL-17-producing T cells, with no formulation eliciting Th2 polarization. Despite similar antibody isotype profiles, rACP+diABZI generates two-fold higher human serum bactericidal activity (hSBA) than rACP+CpG+MPLA and produces antibodies that reverse ACP-mediated inhibition of human neutrophil lysozyme C. Peptide epitope mapping localizes the immunodominant T cell response to the ACP–lysozyme C interaction interface. Passive transfer of immune sera does not recapitulate protection, implicating cellular and mucosal immunity rather than circulating antibodies as the primary effectors. Vaccine-elicited antibodies inhibit Ng adhesion and invasion in three-dimensional (3D) human vaginal and urethral epithelial *in vitro* models. These findings demonstrate that co-induction of T_H_1 and T_H_17 immunity is required for vaccine-mediated protection against Ng, establish diABZI as the first mucosal vaccine adjuvant effective against a bacterial pathogen, and identify ACP as a protective gonococcal antigen.

## RESULTS

### Mucosal rACP+diABZI vaccination accelerates clearance of experimental gonococcal infection

We tested whether immunization with recombinant Ng ACP (rACP) confers protection against gonococcal challenge in female C57BL/6 mice across three independent cohorts that paired rACP with no adjuvant, with CpG+MPLA, or with the STING agonist diABZI (Supplementary Fig. 1). Mice received four intranasal immunizations two weeks apart or, for the rACP+CpG+MPLA group, an intramuscular prime followed by three intranasal boosts. Animals were then challenged intravaginally with Ng strain WHO X (H041), the first isolate identified with high-level resistance to ceftriaxone, cefixime, and other antibiotic classes ^34,35^. It belongs to the ST7363 lineage, which circulates globally across both urogenital and extragenital sites ^34,36^, making it a clinically relevant challenge strain for vaccine evaluation. Vaginal swabs were collected on Days 1, 3, 5, and 7 post-challenge, and clearance was defined as two consecutive culture-negative timepoints. Mice immunized with rACP cleared infection at the same rate as PBS controls (*p* = 0.62), and overall bacterial burden, measured as bioburden area under the curve (AUC), did not differ between groups (*p* = 0.644; Fig. 1a–c). Mice that received rACP+CpG+MPLA likewise cleared infection at the same rate as adjuvant only control (*p* = 0.74), with no AUC difference (*p* = 0.386; Fig. 1d–f). In contrast, rACP+diABZI immunization yielded accelerated clearance relative to diABZI (*p* = 0.038; Fig. 1g). The median bioburden in rACP+diABZI vaccinated mice fell sharply by Day 3 (61 vs 10⁴ CFU/mL in diABZI controls; ∼170-fold reduction), reached the limit of detection by Day 5 and remained there through Day 7, representing a >7,000-fold reduction relative to diABZI control median of 1.4×10^5^ CFU/mL (Mann-Whitney *p* = 0.00038; Fig. 1h). The seven-day AUC trended lower in rACP+diABZI vaccinated mice but did not reach statistical significance (*p* = 0.138; Fig. 1i). Together, these experiments show that protection against Ng requires both the rACP antigen and diABZI and that neither rACP alone nor rACP in combination with CpG and MPLA was sufficient to elicit protection.

**Figure 1.**
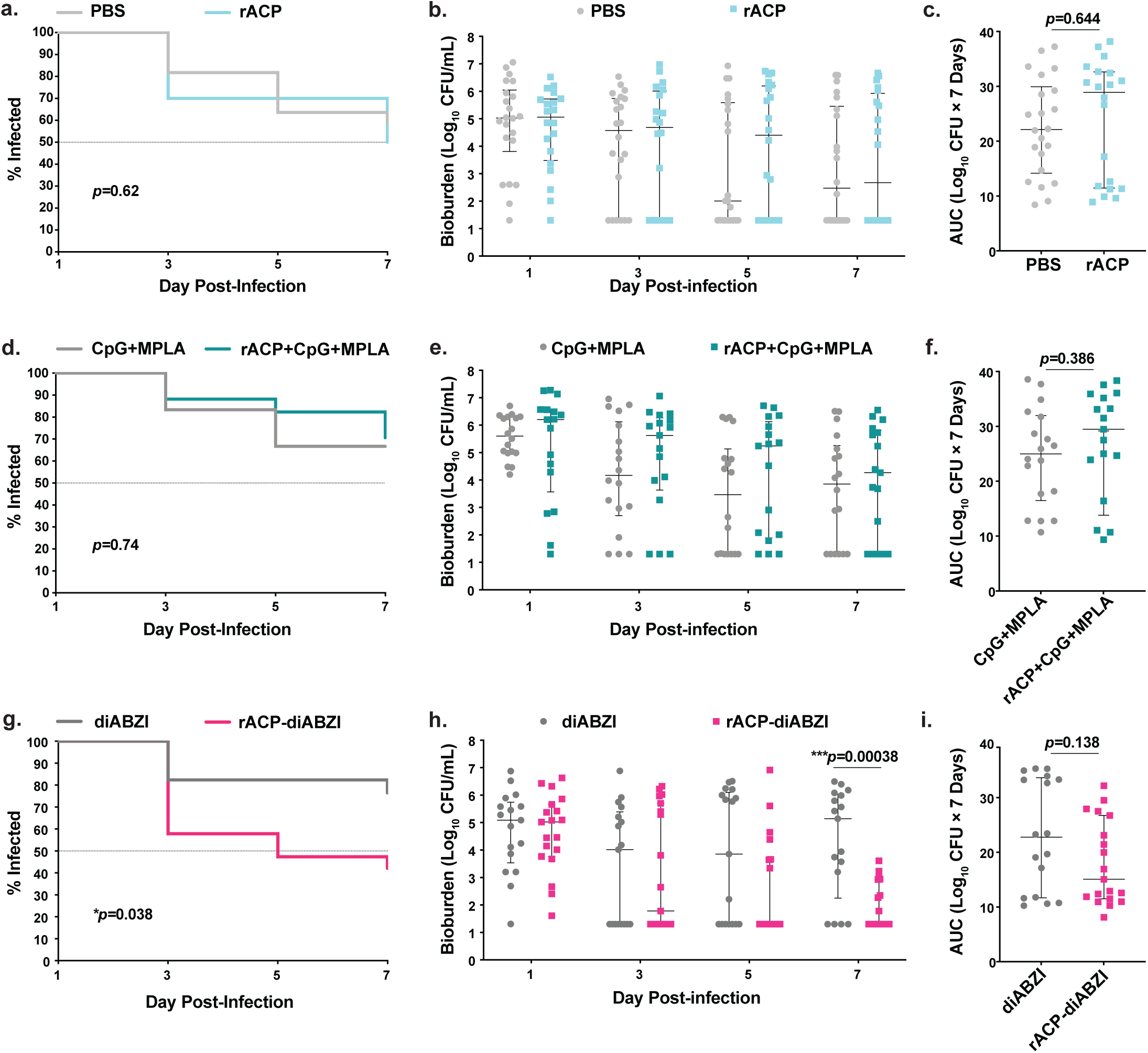
Intranasal rACP+diABZI vaccine accelerates clearance of *Neisseria gonorrhoeae* in the mouse lower genital tract infection model. Immunization–challenge experiments were performed in three independent cohorts of female C57BL/6 mice (*n*=17-22/group): PBS or rACP (**a–c**), CpG+MPLA or rACP+CpG+MPLA (**d–f**), and diABZI or rACP+diABZI (**g–i**). Percentage of mice colonized over time is shown in a, d, and g; bacterial burden in individual mice on days 1, 3, 5, and 7 post-infection in b, e, and h; and cumulative bacterial burden across the 7-day period (AUC, log_10_ CFU×7 days) in c, f, and i. The dashed line in a, d, and g marks 50% colonization. Survival curves were compared by log-rank (Mantel–Cox) test, bacterial burden at each time point by multiple Mann–Whitney tests with Benjamini–Hochberg FDR correction, and AUC by Mann–Whitney test. Bars show median with interquartile range. Source data are provided as a Source Data file.

### rACP+diABZI and rACP+CpG+MPLA induce comparable antigen-specific systemic and mucosal humoral responses

We next examined the specificity of antibody responses by immunoblotting using pooled terminal sera and vaginal lavages probed against purified rACP and whole-cell lysates of Ng FA1090 and its isogenic Δ*acp* mutant (Supplementary Figs. 2-7). As expected, neither systemic or mucosal IgG and IgA from PBS- and adjuvant-treated animals showed reactivity against any antigen, establishing the specificity of the assay (Supplementary Figs. 2a-d and Figs. 3-5). In contrast, serum and vaginal IgG reacted with rACP in all vaccine groups but the detected signal was more apparent in samples derived from mice immunized with adjuvanted vaccines (Supplementary Fig. 2a, c). Additionally, only serum IgG from rACP+diABZI-immunized cohort bound to ACP in the FA1090 wild-type whole cell lysate but not in the Δ*acp* mutant, confirming specificity of the response (Supplementary Fig. 2a). All three vaccine groups produced serum IgA and vaginal IgG that recognized rACP (Supplementary Fig. 2b and c, respectively), and none elicited detectable vaginal IgA that cross-reacted with any of the examined antigens (Supplementary Fig. 2d). To assess cross-strain breadth of the antibodies, we probed pooled sera from each rACP-containing group against whole-cell lysates of FA1090, the Δ*acp* mutant, and 14 contemporary WHO 2016 *Ng* reference strains spanning diverse genetic and geographic backgrounds (Supplementary Figs. 2e-g and 5–7). Serum IgG from rACP-immunized mice cross-reacted only with ACP in WHO P and WHO U, the strains with the highest ACP expression^27^ (Supplementary Figs. 2e and 5), whereas serum IgG from rACP+CpG+MPLA mice cross-reacted with ACP from 13 of 14 strains — all except WHO L at matched exposure (Supplementary Figs. 2f and 6). Signal intensity was consistently higher in the rACP+diABZI IgG sera compared to the two other vaccine groups and cross-reacted with ACP from all WHO 2016 Ng strains (Supplementary Figs. 2f and 7). We next quantified systemic and mucosal antibody responses by ELISA 10 days after the final immunization in sera and in terminal vaginal lavages, respectively (Fig. 2). Across all vaccine formulations the systemic total IgG, IgG1, IgG2c, IgG3 and IgA were significantly increased compared to corresponding control samples (all *p* < 0.001; Figure 2a-c and e, f). Intranasal immunization with rACP alone elicited robust serum total IgG and IgG1 (median log_10_ AUC 4.40 [interquartile range, IQR, 3.40–5.02] and 4.48 [3.16–5.35], respectively) and lower IgG2c (2.69 [0–3.30]), with minimal IgG3 and IgA induction in most mice and no detectable vaginal IgG or IgA (Figure 2). Administration of rACP with adjuvants markedly enhanced all systemic and mucosal Ig responses. Both rACP+CpG+MPLA and rACP+diABZI induced significantly higher serum levels of total IgG, IgG1, and IgG2c compared to rACP alone (all *p* < 0.0001; Fig. 2a–c). IgG2c was particularly responsive to adjuvant as it was only modestly induced by rACP alone (median log_10_ AUC 2.69 [0–3.30]), it rose to 4.84 [4.48–5.21] in rACP+CpG+MPLA and 5.17 [4.95–5.33] in rACP+diABZI mice (Fig 2c). We next assessed IgG subclass induction as well as log_10_(IgG2c/IgG1) ratio as an indicator of T helper (T_H_) polarization. IgG1 predominated in all antigen-containing formulations, consistent with intranasal protein antigen delivery in C57BL/6 mice ^37^. In rACP-only mice, the ratio was strongly IgG1-skewed (median −2.08 [−2.72 to −1.77]), confirming that the four-dose intranasal regimen drives detectable class switching consistent with T_H_2 polarization even in the absence of adjuvant. Adjuvant addition shifted the ratio significantly further toward IgG2c indicating higher T_H_1 polarization (rACP+CpG+MPLA median −1.27 [−1.48 to −1.00], *p* < 0.01; rACP+diABZI median −1.07 [−1.29 to −0.89], *p* < 0.001), with no difference between the two adjuvanted groups (*p* > 0.05), indicating that both formulations promoted comparable T_H_1-associated class switching. Moreover, both rACP+CpG+MPLA and rACP+diABZI induced significantly higher serum IgG3 and IgA than rACP alone (Fig. 2e–h, all *p* < 0.0001). Vaginal antibodies, undetectable in rACP-alone mice, were also robustly induced by both adjuvanted formulations. Vaginal total IgG reached a median log_10_ AUC of 2.52 [IQR 0.89–3.06] in rACP+CpG+MPLA and 2.36 [1.32–2.80] in rACP+diABZI mice, and vaginal IgA reached 1.06 [0–1.89] and 1.28 [0.92–1.96], respectively (all *p* < 0.0003 vs rACP alone). Remarkably, all serum and vaginal antibody titers were comparable between the two adjuvanted groups (all *p* > 0.05). Longitudinal serum and vaginal ELISA across the immunization course confirmed that seroconversion kinetics and IgG and IgA antibody magnitudes were comparable between rACP+CpG+MPLA and rACP+diABZI at every timepoint examined (Supplementary Fig. 8), establishing that this equivalence is not a terminal-sampling artifact. Together, these data suggest that antibody magnitude alone may not account for differences in Ng clearance.

**Figure 2.**
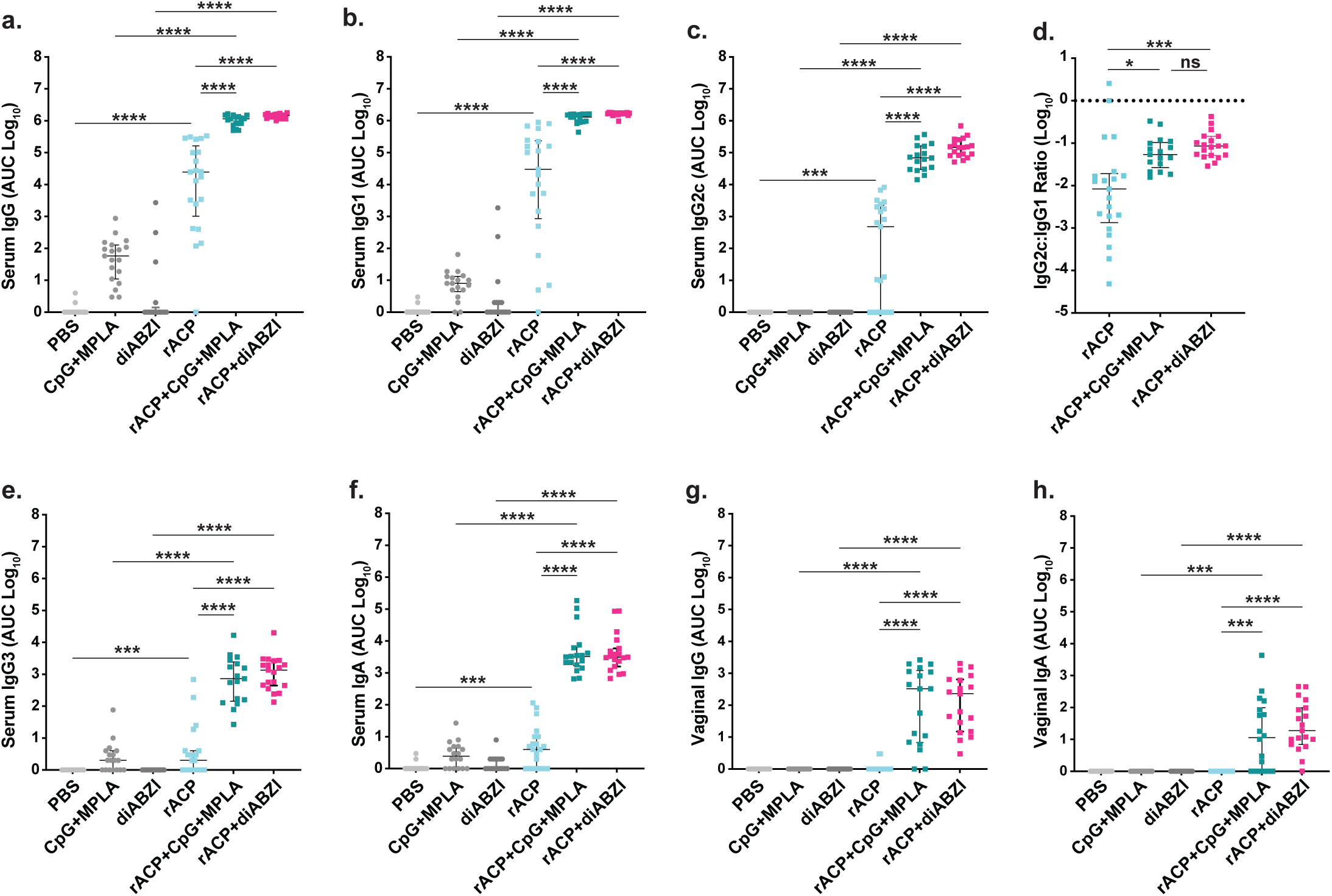
rACP+diABZI and rACP+CpG+MPLA elicit comparable systemic and mucosal antibody responses against rACP. Sera and vaginal lavages collected 10 days after the final boost were assayed by ELISA against rACP; endpoint binding is expressed as area under the dilution curve (AUC) and log_10_-transformed. Serum total IgG (**a**), IgG1 (**b**), IgG2c (**c**), IgG3 (**e**), and IgA (**f**) titers are shown across all six immunization groups, with vaginal IgG (**g**) and IgA (**h**) titers in the same groups. **d**, IgG2c:IgG1 ratio in rACP-containing groups, calculated as log_10_(IgG2c) − log_10_(IgG1); control groups (PBS, CpG+MPLA, diABZI) were excluded because IgG2c was undetectable in their absence, and the dotted horizontal line marks ratio = 0. Comparisons were performed by Kruskal–Wallis test with Dunn’s multiple comparisons post-hoc test and Bonferroni correction. Bars show median with interquartile range. Source data are provided as a Source Data file.

**Figure 3.**
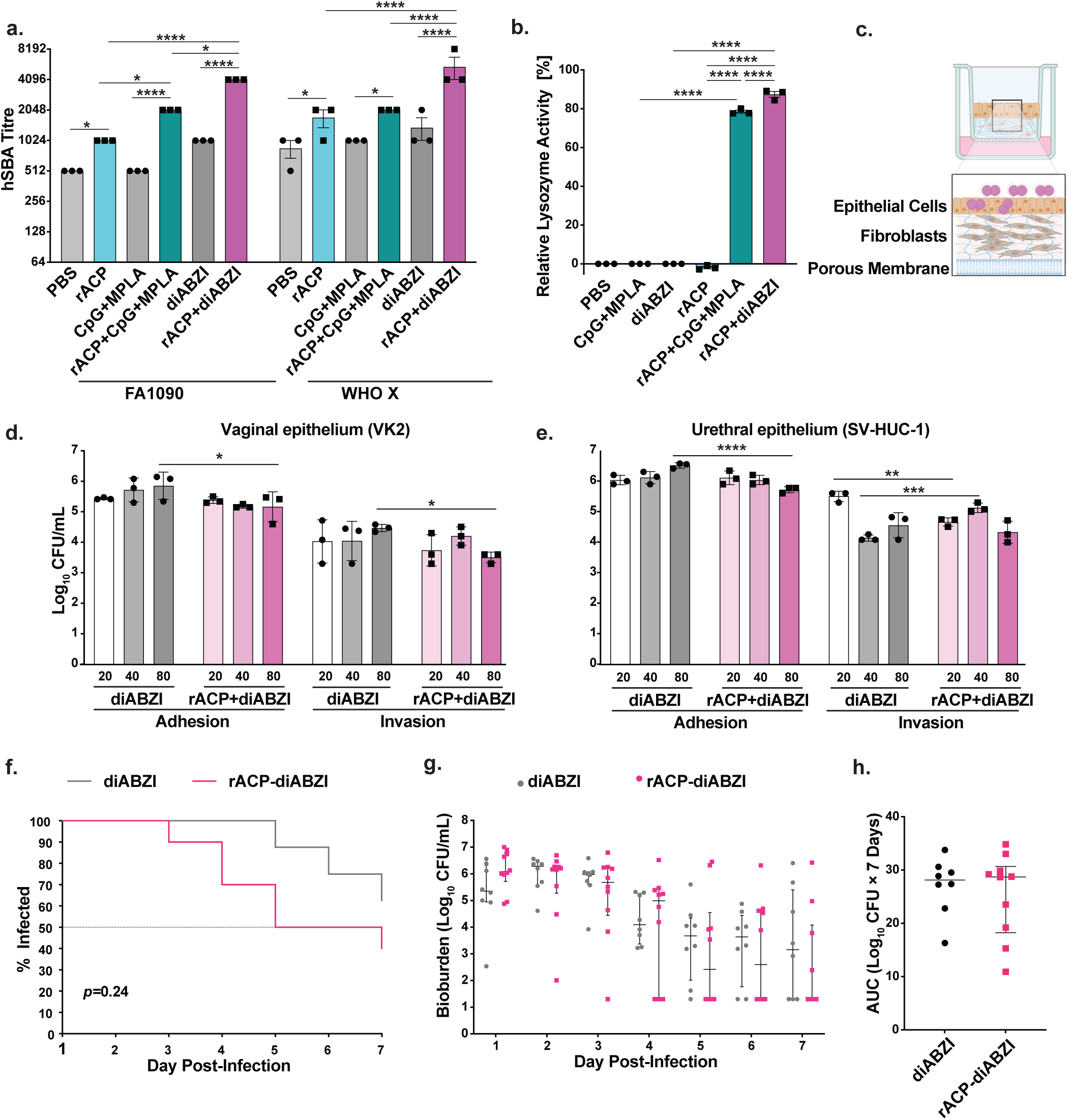
rACP+diABZI elicits the highest serum bactericidal activity, restores lysozyme C function, and generates antibodies that block *N. gonorrhoeae* adhesion and invasion in 3D human epithelial models, but circulating antibodies alone do not confer protection. **a,** Human serum bactericidal activity (hSBA) titers of pooled sera against *N. gonorrhoeae* FA1090 and WHO X. **b,** Relative lysozyme activity rescued by pooled immune sera using human neutrophil-derived lysozyme C, expressed as percent of sham-immunized control. **c,** Schematic of the 3D vaginal/urethral co-culture transwell model. **d, e,** Adhesion and invasion of piliated, Opa⁻ *N. gonorrhoeae* WHO X (MOI 10) on vaginal VK2/E6E7 (**d**) and urethral SV-HUC-1 (**e**) 3D co-cultures, after pre-incubation of bacteria with sera at 1:20, 1:40, or 1:80 dilutions. **f–h,** Passive transfer of pooled, heat-inactivated immune sera into recipient mice prior to intravaginal challenge with *N. gonorrhoeae* WHO X, showing percent infected (f), bacterial burden during the 7-day infection course (**g**), and cumulative bacterial burden (AUC, log_10_ CFU × 7 days; h) hSBA titers were log_2_-transformed and compared by two-way ANOVA (strain × treatment) with Tukey’s multiple comparisons test; lysozyme rescue by one-way ANOVA with Šídák’s multiple comparisons test; adhesion and invasion by two-way ANOVA with Dunnett’s multiple comparisons test. Passive transfer survival was compared by log-rank (Mantel–Cox) test, bacterial burden by multiple Mann–Whitney tests with Benjamini–Hochberg FDR correction, and AUC by Mann–Whitney test. In **a, b, d,** and **e,** *n*=3 independent experiments; bars show mean±SEM. In **g** and **h,** bars show median with interquartile range. Source data are provided as a Source Data file.

### Vaccination with STING agonist elicits significantly elevated human complement-dependent serum bactericidal activity

To assess the functional capacity of vaccine-induced antibodies, we first used human complement-dependent serum bactericidal activity (hSBA) against Ng strains FA1090 and WHO X. PBS and adjuvant-only controls yielded baseline hSBA titers (mean 512–1024 against FA1090; 853–1365 against WHO X; Fig. 3a). Immunization with rACP alone significantly elevated hSBA compared to PBS against both FA1090 (mean 1024±0) and WHO X (mean 1707± 341; *p*≤0.05). Sera from rACP+CpG+MPLA-immunized mice had significantly increased hSBA titers relative to rACP and adjuvant control against FA1090 (mean 2048±0) and compared to CpG+MPLA against the WHO X (mean 2048 ± 0; *p*< 0.0001). Formulating rACP with diABZI led to the highest hSBA titers of any group, reaching mean titers of 4096±0 against FA1090 and 5461±1365 against WHO X, significantly exceeding PBS, rACP alone, CpG+MPLA alone, diABZI alone, and rACP+CpG+MPLA (all *p* < 0.0001 against WHO X; Fig. 3a). As such, despite comparable total antibody titers between the two adjuvanted groups, diABZI induced significantly higher human complement-mediated bactericidal activity compared to CpG+MPLA.

### Immunization with rACP+diABZI induces ACP-neutralizing antibodies that restore innate antimicrobial defense

ACP and SliC are two proteinaceous inhibitors of human lysozyme C encoded by Ng, ^28–31^ yet only immunization with ACP elicits neutralizing antibodies in mice and rabbits that rescue lysozyme activity against bacterial peptidoglycan ^27,38^. We therefore compared whether vaccine-elicited antibodies could neutralize the ACP-mediated lysozyme blocking phenotype. For this, we used recombinant lysozyme C from human neutrophils. Sera from mice immunized with rACP did not rescue lysozyme activity above background compared to PBS, CpG+MPLA, or diABZI treatment (Fig. 3b). In contrast, rACP+CpG+MPLA and rACP+diABZI vaccines elicited serum antibodies that restored lysozyme activity (mean 78.5% and 87.4%, respectively versus corresponding adjuvant-only controls and rACP (*p* < 0.0001). Notably, rACP+diABZI sera restored significantly greater lysozyme activity than rACP+CpG+MPLA sera (mean difference 8.9%, *p* < 0.0001), indicating that diABZI as an adjuvant generates antibodies with higher capacity to block the ACP–lysozyme interaction and restore innate antimicrobial defense.

### rACP-diABZI vaccine elicits antibodies interfering with Ng adhesion and invasion in human 3D epithelial culture models

Meningococcal ACP functions as both an adhesin and a minor invasin. ACP-expressing bacteria demonstrate robust association with epithelial, endothelial, and meningeal cells in conventional two-dimensional culture, and anti-ACP sera can block this interaction in an inoculum-dependent manner ^28^. However, two-dimensional monolayer systems do not recapitulate the structural complexity, epithelial barrier integrity, or transport dynamics of intact mucosal tissue, limiting the physiological relevance of adhesion and invasion assays conducted in these simplified environments ^39^. To address whether rACP-diABZI-induced antibodies interfere with gonococcal adhesion and invasion at mucosal surfaces approximating the site of infection, we developed a transwell-based 3D model with vaginal (VK2/E6E7) or urethral (SV-HUC-1) epithelial cells co-cultured with primary fibroblasts (HUtF) within a GelMA hydrogel, which demonstrated support for epithelial organization and barrier function in gynecologic and mucosal tissue applications ^40,41^ (Fig. 3c). We first assessed epithelial barrier integrity in the 3D models prior to Ng infection. Stable transepithelial electrical resistance (TEER) values were acquired after 7 days of co-culture for both cell lines, with a significant increase in barrier integrity observed on day 7 (Supplementary Fig. 9). SV-HUC-1 cells exhibited slightly higher TEER values at this time point, but no significant differences were observed between cell lines. With vaginal epithelial cells, sera from rACP+diABZI-vaccinated mice reduced both adhesion and invasion of Ng WHO X (Pili+, Opa−) relative to sera from diABZI only mice in a dilution-dependent manner, with significant reductions at the 1:80 dilution for both adhesion (*p* = 0.02) and invasion (*p* = 0.02; Fig. 3d). In urethral epithelial cells, adhesion was similarly reduced in the presence of rACP+diABZI sera, with a significant decrease at the 1:80 dilution (*p* < 0.0001; Fig. 3e). Invasion in the urethral model showed a more complex dose–response pattern. A significant reduction in bacterial invasion was observed at the 1:20 dilution (*p*=0.002) rather than at 1:80, as in the vaginal model. Across both tissue models, sera from rACP+diABZI-vaccinated mice consistently reduced Ng adhesion and invasion compared with diABZI-only sera, demonstrating that vaccine-induced antigen-reactive antibodies interfere with gonococcal interaction with both female and male mucosal epithelium.

### Passive transfer of immune sera is insufficient to confer protection

To determine whether circulating antibodies alone are sufficient for protection, we performed passive immunization by transferring pooled sera from diABZI-only or rACP+diABZI-vaccinated mice prior to Ng challenge. Mice receiving rACP+diABZI sera showed a trend toward accelerated clearance relative to diABZI sera recipients, but this did not reach statistical significance (logrank *p*=0.24; Fig. 3f). Daily bioburden (Fig. 3g) and cumulative bacterial load over seven days (AUC *p*=0.138; Fig. 3h) were also not significantly different between groups, although point estimates trended lower in the rACP+diABZI group. The failure of passive transfer to recapitulate the protection observed with active immunization suggests that circulating antibodies are not primary mediators of vaccine-induced clearance and implies cellular immunity and/or locally generated mucosal immune responses as essential components of diABZI-adjuvanted ACP-mediated protection.

### diABZI uniquely drives Th1/Th17 co-polarization

To further elucidate the mechanisms of STING agonist-associated clearance and decrease in Ng bioburden, we aimed to characterize T helper responses (Fig. 4, Supplementary Figs. 10-14). For this splenocytes from immunized mice were stimulated *ex vivo* with a pool of overlapping 15-mer peptides spanning full-length ACP and assessed by ELISpot for IFNγ (Th1), IL-4 (Th2), and IL-17 (Th17) production (Fig. 4a-d). Both rACP+diABZI and rACP+CpG+MPLA induced comparable numbers of IFNγ-producing cells (mean 361 and 357 spots, respectively, *p* = 0.9999 between groups). These were both highly significant compared to rACP-only vaccination (mean 40.5) and adjuvant-only controls (*p*<0.0001, Fig. 4a). To assess MHC-I-restricted responses, DC2.4 cells stably transduced with a lentivector encoding ACP ^42^ were co-cultured with splenocytes from vaccinated mice. Adjuvanted vaccines induced IFNγ secretion; however, rACP+CpG+MPLA elicited significantly more IFNγ-producing cells than rACP+diABZI (mean 350.3 vs. 251.4 spots; *p* = 0.0087; Fig. 4b), suggesting that CpG+MPLA may lead to more effective priming of antigen-specific CD8+T cells. No formulation induced detectable IL-4-secreting cells (Fig. 4c), confirming the absence of T_H_2 polarization of CD4+ T cells across all groups. Intriguingly, only rACP+diABZI induced significant IL-17 production (mean 229 spots; *p* < 0.0001; Fig. 4d) and rACP+CpG+MPLA did not elicit IL-17 above background (mean 5.1). Thus, while both adjuvants drive T_H_1 (IFNγ) responses, only diABZI co-induces T_H_17, a distinction that parallels the divergent vaccine-mediated protection outcomes observed in the challenge experiments.

**Figure 4.**
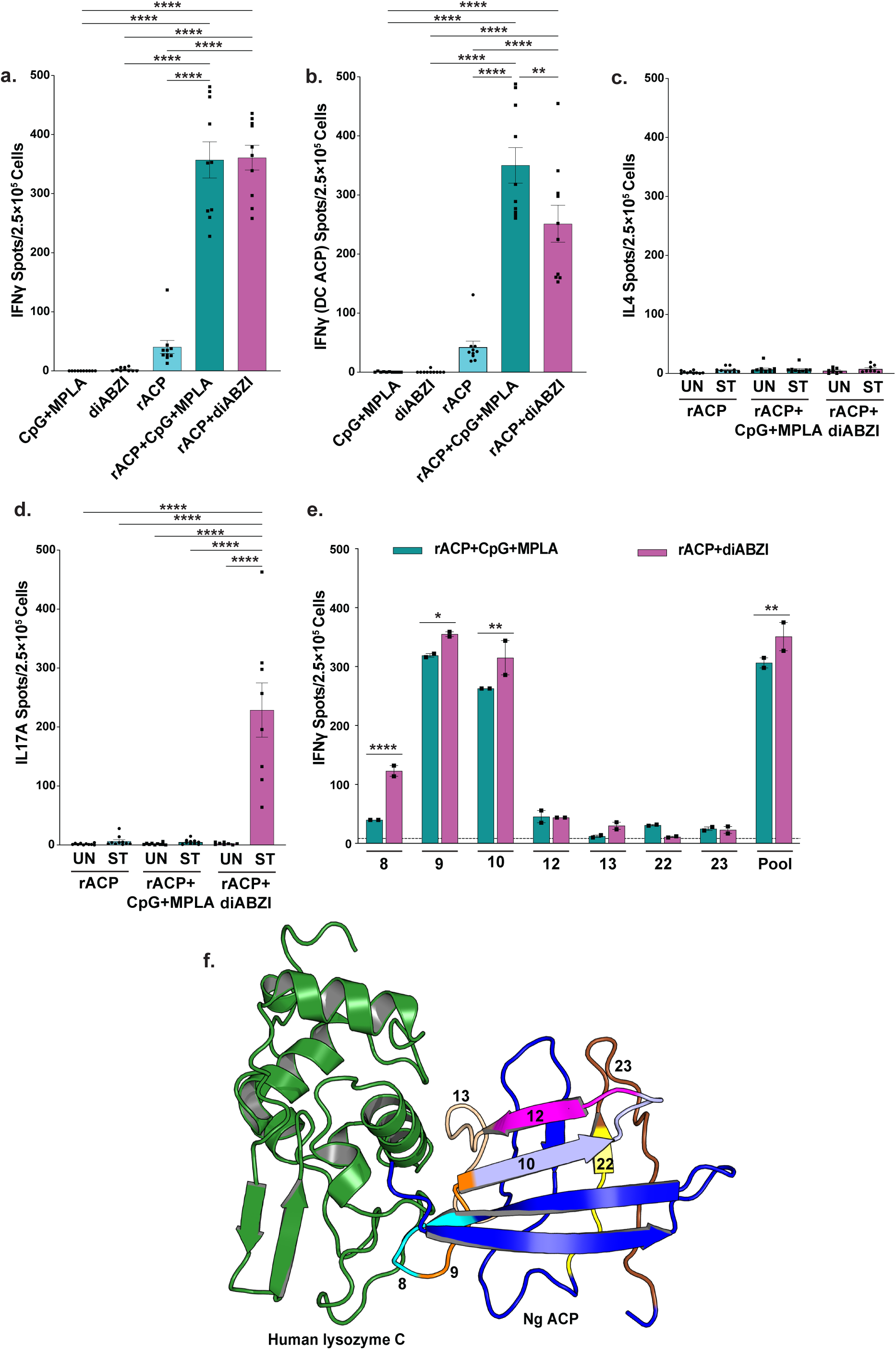
rACP+diABZI uniquely co-induces T_H_1 and T_H_17 responses; immunodominant ACP epitopes localize to the human lysozyme C interface. Splenocytes from immunized mice were assayed by ELISpot following co-culture with DC2.4 cells at a 1:10 (DC2.4:splenocyte) ratio; spot counts in panels **a–e** are normalized to 2.5×10^5^ cells. (a) IFNγ-secreting cells after stimulation with the 23-peptide ACP pool. (b) IFNγ-secreting cells after co-culture with DC2.4 cells stably transduced with a lentivector encoding ACP, assaying MHC class I–restricted CD8+ T cell responses. Panels a and b show all five immunization groups (CpG+MPLA, diABZI, rACP, rACP+CpG+MPLA, rACP+diABZI; *n*=10 mice/group, individual values plotted). (c) IL-4–secreting and (d) IL-17A–secreting cells in unstimulated (UN) and ACP peptide pool–stimulated (ST) wells; only the three rACP-containing groups are shown (*n*=10/group). (e) IFNγ responses to individual ACP peptides for splenocytes pooled within each adjuvanted group and plated in technical duplicate; data are shown for peptides 8–10, 12, 13, 22, 23, and the full pool. The dotted horizontal line marks the immunodominance threshold (mean+2SD of non-reactive peptides); peptides 8 (FNKQGLTTYASAVIN), 9 (GLTTYASAVINGKRV), and 10 (YASAVINGKRVQMPI) exceeded this threshold. (f) ClusPro docking of human c-type lysozyme (PDB 7XF6, green) and *N. gonorrhoeae* ACP (PDB 6GQ4, blue), with the seven peptides screened in (e) highlighted in color and labelled (PyMOL 3.1). Bars show mean ± SEM. Comparisons in a–d were performed by Kruskal–Wallis test with Dunn’s multiple comparisons post-hoc test and Bonferroni correction. In (e), within-peptide comparisons of rACP+diABZI versus rACP+CpG+MPLA were derived from two-way ANOVA with Šídák’s multiple comparisons test; \**p* < 0.05, \*\**p* < 0.01, \*\*\**p* < 0.001, \*\*\*\**p* < 0.0001. Source data are provided as a Source Data file.

### Immunodominant ACP epitopes localize to the lysozyme C interaction interface

To map immunodominant T cell epitopes of the ACP protein, we pooled splenocytes from each vaccine cohort, singly stimulated them ex vivo with each of the 23 ACP-derived 15-mer peptides, and performed IFNγ ELISpot (Fig. 4e, Supplementary Fig. 10). Three peptides were found to induce IFNγ secretion: peptide 9 (GLTTYASAVINGKRV; 355 and 319 spots per 2.5×10^5^ cells for rACP+diABZI and rACP+CpG+MPLA, respectively), peptide 10 (YASAVINGKRVQMPI; 315 and 263), and peptide 8 (FNKQGLTTYASAVIN; 123 and 40). Two-way ANOVA across these peptides confirmed a significant adjuvant effect (*F*_1,16_ = 24.6, *p* = 0.0014) and a significant peptide×adjuvant interaction (*F*_7,16_=5.16, *p* = 0.003), indicating that the diABZI advantage was peptide-specific rather than uniform. Peptide 8 stood out as the most adjuvant-discriminating epitope: rACP+diABZI elicited an ∼3-fold higher response than rACP+CpG+MPLA (*p* < 0.0001), the largest within-peptide difference in the dataset. Smaller but significant STING agonist-driven advantages were also seen at peptides 9 (*p* = 0.02) and 10 (*p* = 0.003) and the full peptide pool (*p* = 0.009). The remaining peptides — 12, 13, 22, and 23 — produced responses at or near the assay background (dashed line, Fig. 4e), with no peptide exceeding ∼45 spots in either group. The three immunodominant peptides share an overlapping core within the FNKQGLTTYASAVINGKRVQMPI region of ACP. Protein–protein docking of human lysozyme C and ACP revealed a stable complex with ACP positioned along the lysozyme C surface (Fig. 4f), and all three immunodominant peptides (8, 9, and 10) mapped to the lysozyme C–ACP binding interface while remaining surface-exposed within the complex. Strikingly, peptide 8 localized to the β-strand at the heart of the lysozyme C–ACP contact, placing the most diABZI-responsive T cell epitope directly at the inhibitory interface.

### Integrated analysis identifies hSBA as the sole immune correlate of accelerated Ng clearance

To identify immune readouts associated with accelerated clearance after rACP+diABZI immunization, we analyzed nine parameters in the 19 vaccinated mice: serum IgG, IgG1, IgG2c, IgG3, and IgA; vaginal IgG and IgA; serum lysozyme rescue activity; and serum bactericidal activity (hSBA) against the challenge strain WHO X (Figure 5, Supplementary Table 1 and Supplementary Figs. 15-16). Univariate Cox proportional hazards regression for time to clearance identified hSBA titer as the strongest predictor of accelerated clearance (*p* < 0.001), with serum IgA as the only other significant association (HR=1.91 [95% CI 1.09–3.37], *p*=0.025). Serum IgG, IgG1, IgG2c, IgG3, vaginal IgG, vaginal IgA, and lysozyme rescue activity showed no association with clearance (all *p*≥0.12; Fig. 5a). Day-resolved Spearman correlation matrices between each immune parameter and bioburden (Supplementary Fig. 16) identified hSBA as the only immune parameter significantly correlated with bioburden, with peak negative correlation at Day 3 (r = −0.53). Stratifying animals at hSBA = 2,048 (the threshold maximizing separation between clearance groups) revealed an obvious dichotomy in clearance kinetics: 9 of 11 mice with hSBA ≥2,048 cleared infection by Day 7, compared with 2 of 8 mice with hSBA <2,048 (log-rank χ² = 5.8, *p* = 0.02; Fig. 5b). The high-titer group cleared rapidly, with most animals clearing by Day 3, whereas the majority of low-titer animals failed to clear by the end of the experiment. ROC analysis of Day 3 clearance reinforced the hierarchy seen in the Cox regression. hSBA titer discriminated Day 3 clearers from non-clearers nearly perfectly (AUC = 0.95), while no other readout exceeded AUC 0.72. Serum IgG2c (AUC = 0.51), vaginal IgA (AUC = 0.49), and serum IgG1 (AUC = 0.48) performed at chance (Fig. 5c). Plotting individual hSBA titers across the four clearance categories revealed a monotonic relationship between titer magnitude and clearance speed (Fig. 5d). Mice that cleared by Day 3 had a median hSBA titer of 4,096, four-fold higher than mice that never cleared (median 1,024; Day 3 vs. Never, *p*=0.0023; Kruskal–Wallis across all four clearance categories, *p*=0.0061). All eight Day 3 clearers had hSBA titers at or above 2,048, while only one of eight never-cleared animals exceeded that threshold. To visualize the joint relationship between pre-challenge titer, bioburden, and clearance trajectory at the per-animal level, we plotted all 19 mice across Days 1, 3, 5, and 7 post-challenge, ordered by hSBA titer (highest at top) with bubble size proportional to log10 CFU/mL and color encoding clearance day (Fig. 5e). The integrated view mirrors the statistical findings: high-titer animals at the top of the plot showed rapid bioburden attenuation and predominantly cleared by Day 3, whereas low-titer animals at the bottom retained measurable bioburden through Day 7 and largely failed to clear. The transition between these regiments tracked closely with the hSBA = 2,048 threshold identified in panels b–d. Together, these analyses (univariate Cox regression, threshold-stratified Kaplan–Meier estimation, ROC discrimination, categorical comparison across clearance times, and integrated longitudinal visualization) converge on hSBA titer against WHO X as the dominant correlate of accelerated clearance in rACP+diABZI-immunized mice. Serum IgA reached significance in Cox regression but performed only modestly in ROC (AUC = 0.67) and did not separate clearance categories, placing it below hSBA in predictive value. We therefore identify hSBA titer as a candidate functional correlate of STING agonist diABZI+rACPvaccine-induced protection in the murine lower genital tract Ng model and a quantitative readout to anchor downstream antigen and adjuvant comparisons.

**Figure 5.**
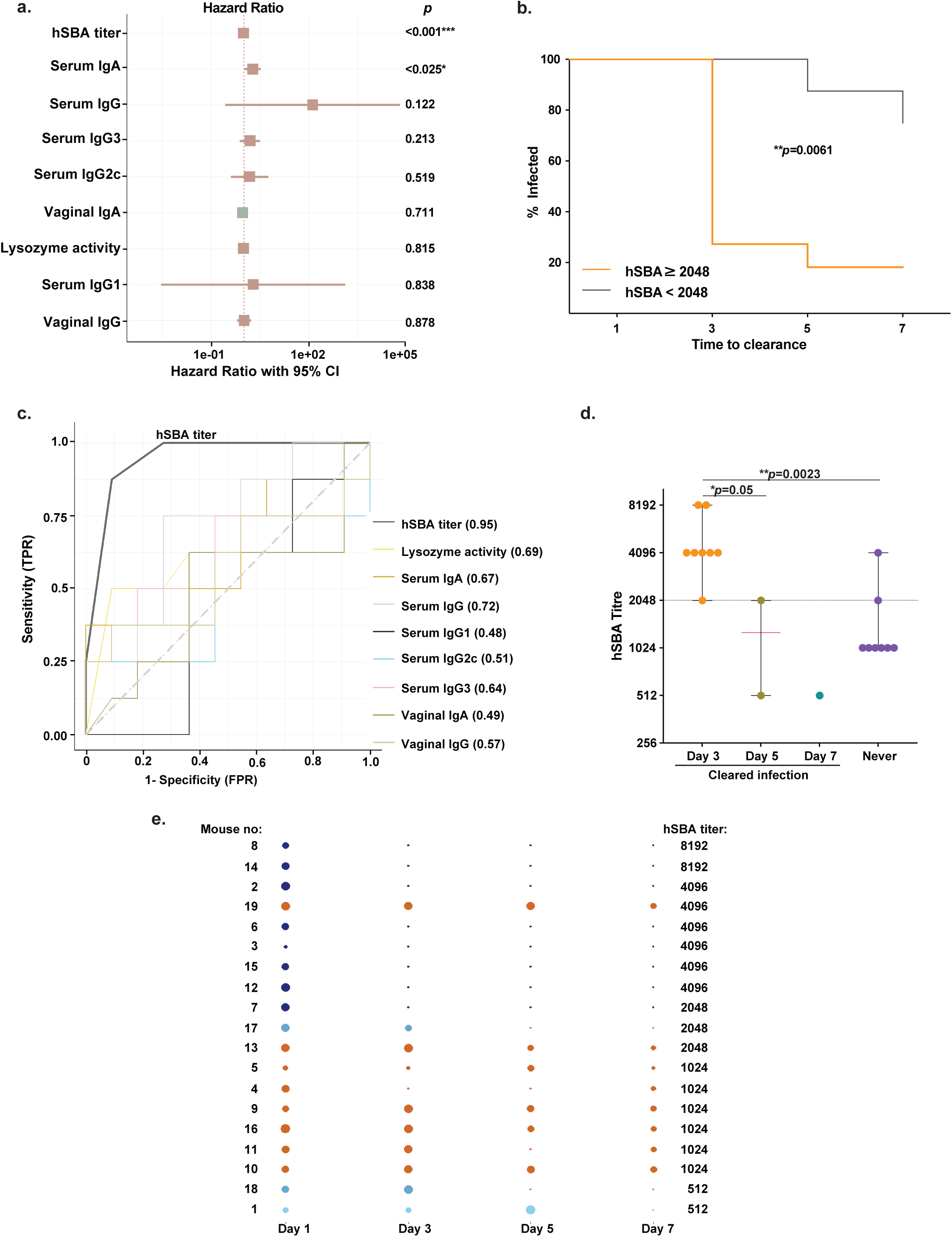
hSBA titer against the challenge strain is the dominant correlate of accelerated *Neisseria gonorrhoeae* clearance after mucosal rACP+diABZI immunization. Pre-challenge serum and vaginal samples from 19 mice immunized with rACP+diABZI were analyzed for nine immune readouts and tested for association with time to bacterial clearance after intravaginal challenge with Ng strain WHO X. (a) Forest plot of hazard ratios from univariate Cox proportional hazards regression for time to clearance. Squares indicate point estimates and horizontal lines show 95% confidence intervals; *p*-values from the Wald test are shown on the right. \**p* < 0.05; \*\*\**p* < 0.001. (b) Kaplan–Meier estimate of percent infected over time for animals stratified at the hSBA = 2,048 threshold (orange, hSBA ≥2,048, *n*=11; black, hSBA <2,048, *n*=8). *p*-value from *log*-rank test. (c) Receiver-operating characteristic (ROC) curves discriminating Day 3 clearers from non-clearers for each of the nine immune readouts; AUC values are shown in the legend. Diagonal indicates chance performance. (d) Pre-challenge hSBA titers stratified by day of bacterial clearance (Day 3, *n* = 8; Day 5, *n* = 2; Day 7, *n* = 1; Never cleared, *n* = 8). Horizontal bars indicate medians and interquartile ranges. Pairwise comparisons by Mann–Whitney U test; overall comparison across the four categories by Kruskal–Wallis test. \**p* < 0.05; \*\**p* < 0.01. (e) Per-animal longitudinal bubble plot of all 19 vaccinated mice, sorted on the y-axis by pre-challenge hSBA titer (highest at top). Bubble size is proportional to log_10_ CFU/mL bioburden at Days 1, 3, 5, and 7 post-challenge. Color encodes the day on which each animal cleared infection (dark blue, Day 3; medium blue, Day 5; light blue, Day 7; orange, never cleared). Pre-challenge hSBA titers are listed on the right.

## DISCUSSION

The development of an effective gonorrhea vaccine remains an urgent unmet clinical need. Two vaccines composed of killed whole Ng cells and purified pilin failed in clinical trials decades ago, illustrating the difficulty Ng poses for conventional vaccine design. Longstanding barriers such as antigenic variability, sophisticated host immune evasion strategies, and the absence of established correlates of protection ^19^ have recently been compounded by independent randomized controlled trials that have largely challenged the hypothesis of 4CMenB-mediated cross-protection. The ANRS 174 DOXYVAC trial reported a non-significant 22% relative reduction in gonorrhea incidence (adjusted hazard ratio 0.78, *p* = 0.061) ^18^; MenGO, a non-significant incidence rate ratio of 0.78 (95% CI 0.40–1.51, *p* = 0.457)^17^; and GoGoVax, conducted in gay, bisexual, and other men who have sex with men, reported an overall vaccine efficacy of −0.5% over 24 months of follow-up ^15,16,17^. Exacerbating these cross-protection setbacks, the only recent Ng-specific clinical-stage candidate [GMMA-based engineered OMV formulation (NgG, GSK)] was discontinued by its developer before efficacy data were reported ^1,20^, leaving no active gonorrhea-targeted vaccine in clinical development.

Against this background, we report four advances. First, we identify ACP as a protective gonococcal vaccine antigen in an active immunization–challenge model, the first such demonstration for this conserved dual-function protein. Second, we establish diABZI as the first non-nucleotide STING agonist to elicit protection against a bacterial pathogen. Third, comparison of rACP+diABZI with rACP+CpG+MPLA identifies T_H_1/T_H_17 co-induction as the immunological signature of protection for ACP. Fourth, we demonstrate vaccine-elicited blocking of Ng WHO X adhesion and invasion in a barrier-competent, transwell-based 3D co-culture of human vaginal (VK2/E6E7) and urethral (SV-HUC-1) epithelial cells. This is to our knowledge the first side-by-side evaluation of vaccine-mediated functional blocking in male and female human 3D mucosal models, advancing the field beyond the cervical monolayer (ME-180) systems typically used in gonococcal vaccine studies. Additional observations contextualize these findings within a mechanistic and translational framework: convergence of immunodominant T cell epitopes on the ACP–lysozyme C interface and identification of hSBA as the dominant correlate of accelerated clearance.

T_H_1 polarization has been associated with accelerated gonococcal clearance in protein subunit vaccines LOS 2C7 epitope mimic and Ng OMVs^19,43–46^. For ACP, however, T_H_1 polarization alone was not sufficient for protection. rACP+CpG+MPLA and rACP+diABZI drove equivalent IgG2c class switching, comparable ACP-specific IFNγ responses, and similar serum and vaginal antibody titers, yet only rACP+diABZI co-induced IL-17A, and only rACP+diABZI accelerated clearance. The protective signature for ACP therefore appears to require T_H_17 co-induction in addition to T_H_1 response, an antigen-specific dependence consistent with evidence from recent mucosal gonococcal vaccine studies. Intranasal immunization with MlaA adjuvanted with cholera toxin B subunit (CTB) induced T_H_1, T_H_2, and T_H_17 and high hSBA titers, blocked Ng association with ME-180 epithelial cells, and accelerated clearance in the murine vaginal infection model^47^. A trivalent App+NHBA+MetQ CTB-adjuvanted vaccine administered intranasally^48^ also induced T_H_1/T_H_17 co-polarization and accelerated Ng clearance, and a recent study of the conserved Slam antigen reported a T_H_17-skewed cellular response together with bactericidal and adhesion-blocking antibodies^49^. In parallel, a comparative immunoprofiling study of 4CMenB and Ng OMVs identified polymorphonuclear neutrophil (PMN) influx to the murine genital tract as the immune readout most strongly associated with reduced Ng bioburden^50^ which is mechanistically consistent with T_H_17-driven, IL-17A–dependent neutrophil recruitment. Together, these studies indicate that T_H_17 co-induction broadens the antigen and platform compatibility of protective gonococcal vaccines. This also aligns with vaccine-induced T_H_1/T_H_17 co-protection established for other mucosal and intracellular pathogens, including another sexually transmitted bacterium, *C. trachomatis*^23,24,51^.

Among immune readouts evaluated in individual rACP+diABZI-immunized mice, hSBA titre emerged as the sole robust immune correlate with time to clearance; stratification at hSBA ≥2,048 separated rapid clearers from non-clearers. The female genital tract is an IgG-dominant mucosal compartment^52,53^, and serum IgG is efficiently transported to the genital lumen by the neonatal Fc receptor (FcRn)^54^, providing a mechanistic basis for the linkage between systemic hSBA and mucosal protection. hSBA is the accepted surrogate of protection for *N. meningitidis*, and its predictive value for Ng is under active investigation. Prior *in vivo* evidence is limited to passive immunization with the 2C7 LOS monoclonal antibody^46^ and to C9–/– mouse studies of the TMCP2/GLA-SE LOS mimotope vaccine, in which loss of terminal complement abolished protection despite intact anti-LOS IgG titers, nominating SBA as a correlate of protection^55^. Our per-animal, multi-readout analysis is the first integrated demonstration in an active immunization–challenge model with a defined recombinant antigen, where hSBA dominates other antibody and functional readouts in predicting time to clearance. By contrast, McKeand *et al*. did not detect an hSBA–clearance correlation in their immunoprofiling of 4CMenB and Ng OMVs^50^; and the MenGO trial reported a statistically significant but quantitatively modest hSBA increase after 4CMenB vaccination that did not translate to clinical protection ^17^. We interpret these findings as complementary and reflective of platform-dependent indicators of protection.

An important feature of our findings is the convergence of cellular and humoral immunity on ACP function as an inhibitor of human lysozyme C and as an adhesin. The three immunodominant T cell epitopes (peptides 8, 9, and 10) span the FNKQGLTTYASAVINGKRVQMPI region of ACP, with peptide 8 (FNKQGLTTYASAVIN) lying at the β-strand central to the ACP-lysozyme C contact and showing the largest diABZI-dependent advantage. Antibodies elicited by rACP+diABZI restored human neutrophil lysozyme C activity to 87% of control, significantly above rACP+CpG+MPLA at 78%, demonstrating that the polyclonal antibody response functionally relieves ACP-mediated inhibition of lysozyme C. These antibodies also reduced Ng WHO X adhesion and invasion in human 3D mucosal models, thereby abolishing both the lysozyme-inhibitor and adhesin functions of ACP. Although we did not map B cell epitopes in this study, this dual functional neutralization is consistent with antibodies engaging determinants that overlap or are proximal to the lysozyme-binding and host-contact surfaces of ACP. diABZI-driven T cell help and antibody maturation therefore act in concert to disable two conserved virulence functions of ACP, simultaneously priming cellular effectors against ACP-presenting cells and generating antibodies that block bacterial attachment and relieve ACP-mediated inhibition of an innate antimicrobial defense. Dual engagement of this conserved functional axis offers a mechanistic rationale for why ACP is well suited to a vaccine strategy, and accounts for the broad cross-reactivity of rACP+diABZI sera with ACP from all 14 WHO 2016 reference strains.

## METHODS

### Bacterial strains and growth conditions

The study utilized *Neisseria gonorrhoeae* (Ng) strain FA1090 and the 2016 World Health Organization (WHO) reference panel of Ng isolates ^35,56^. Bacteria were revived from frozen stocks and cultured on gonococcal base (GCB; Difco) agar supplemented with Kellogg’s supplements I (1:100) and II (1:1,000), or on chocolate agar plates, as appropriate for specific assays. Cultures were incubated for 20 hours at 37°C in a humidified atmosphere containing 5% CO_2_ ^57^. Transparent, non-piliated colonies were selected for human serum bactericidal assays (hSBA), whereas piliated bacteria were used in challenge experiments and adhesion/invasion assays. *Escherichia coli* BL21(DE3) carrying pET22M-rACP was maintained on Luria–Bertani (LB) agar and cultured in LB broth supplemented with carbenicillin and kanamycin (50 µg/mL each) ^27^.

### Antigen expression and purification

Recombinant Adhesin Complex Protein (rACP) was overexpressed in *E.coli* BL21(DE3) carrying pET22M-rACP, grown in 8 L of LB medium and purified as previously described ^27^. Bacteria were cultured in an orbital shaker at 37°C and 200 rpm to an OD_600_∼0.7, at which point rACP expression was induced with 1mM isopropyl-β-D-1-thiogalactopyranoside. *E. coli* cells were harvested 4 h after induction by centrifugation at 5,000×*g* for 15 min at 4°C. Pellets were resuspended in binding buffer (50 mM Na_2_HPO4, 150 mM NaCl, 10 mM imidazole, 5 % glycerol, pH 7.5) supplemented with 70 µM lysozyme, 0.1 µM DNase and protease inhibitor (Pierce), and incubated on ice for 30 min. Cells were lysed by French Press at 1,500 psi and centrifuged at 12,000×*g* for 30 min at 4°C. The supernatant was filtered using a 0.45 µm nylon membrane and applied to a HisTrap FF 5 mL Nickel-NTA IMAC column (Cytiva Life Sciences) using an NGC Medium-Pressure Liquid Chromatography system (Bio-Rad). Bound protein was eluted with a 20% gradient of elution buffer (binding buffer containing 250 mM imidazole), pooled, and dialyzed overnight against binding buffer at 4°C using snakeskin tubing (3.5 MWCO) with gentle stirring. To remove the 10His×Tag, purified rACP was incubated with Tobacco Etch Virus (TEV) protease in the presence of 1 mM Ethylenediaminetetraacetic acid (EDTA) and 0.5 mM Dithiothreitol (DTT) at a 1:75 protease:protein ratio overnight at 4°C. TEV protease and cleaved 10His×Tag were removed by a second IMAC pass; the tag-free rACP in the flowthrough was concentrated using a Vivaspin 20 centrifugal concentrator (Sartorius). Protein concentration was determined by the Qubit Protein Assay on a Qubit 4 Fluorometer (Invitrogen), and the purity was assessed by SDS-PAGE and Western blotting.

### Vaccine formulation

Stocks of VacciGrade™ CpG (ODN2395; 10 mg/mL; InvivoGen) and MPLA (10 mg/mL; InvivoGen) were prepared by resuspension in endotoxin-free physiological saline. diABZI (1 mg/mL) was solubilized in endotoxin-free, apyrogenic sterile water. Adjuvants were combined with the appropriate amount of rACP immediately prior to immunization.

### Immunization and challenge studies

The study is reported in accordance with ARRIVE guidelines. Female C57BL/6 SPF mice (4-6 weeks old; Charles River Laboratories) were used in three independent active immunization/challenge experiments evaluating rACP-based vaccine formulations. In the first cohort (*n* = 21-22/group), mice received four intranasal immunizations every two weeks with either endotoxin-free PBS or 30 μg rACP (total volume 10 μL; 5 µL into each nare). In the second cohort (*n* = 17-18/group) mice were primed intramuscularly with 50 µg of rACP formulated with 15 µg of CpG and 10 µg of MPLA (total volume 30 µL), followed by three intranasal boosts every 2 weeks with 20 µg of rACP formulated with 15 µg of CpG and 10 µg of MPLA (total volume 10 µL; 5 µL per nare). The third cohort (*n*= 17-19) received four intranasal doses at two-week intervals with either 5 µg diABZI alone or 30 µg rACP formulated with 5 µg diABZI (total volume 10 µL; 5 µL per nare). Blood was collected via the saphenous vein 10 days after each immunization beginning with the first boost, and vaginal lavages were collected after the first and second boosts and at study completion. Two weeks after the final immunization, vaginal lavages were examined microscopically to determine estrous cycle stage ^58,59^. Mice in their diestrus were injected subcutaneously with 0.5 mg water-soluble β-estradiol (Premarin; Pfizer) on days -2, 0, and +2 relative to Ng infection ^46^. To suppress commensal flora and facilitate Ng colonization, mice received intraperitoneal injections of streptomycin sulfate (2.4 mg; VWR) and vancomycin hydrochloride (0.4 mg; VWR) for four consecutive days beginning two days prior to challenge, and trimethoprim (0.4 mg/mL; MP Biomedicals) was provided in the drinking water ^59^. From day +2 post challenge of streptomycin sulphate (2.4 mg/mL) was added to the drinking water alongside trimethoprim until study completion. Mice were inoculated intravaginally with 10^6^ CFU of piliated Ng WHO X as described ^60^. Vaginal swabs for CFU enumeration were collected using sterile rayon-tipped applicators (Puritan 25-800 1PD 50) on days 1, 3, 5, and 7 post-infection. Swabs were vortexed in GC broth containing 0.05 % (w/v) saponin, and dilutions were plated using an Eddy Jet 2W Spiral Plater (IUL Instruments) on GC agar supplemented with VCNT Inhibitor (vancomycin, colistin, nystatin, trimethoprim sulfate, and streptomycin sulfate; Beckton Dickinson). Plates were incubated for 48 h in 37°C with 7% CO_2_ and imaged on an automated colony counter (SphereFlash; Neutec Group Inc). The lower limit of detection was 20 CFU/mL. Clearance was defined as two consecutive culture-negative timepoints (≤20 CFU/mL, LOD = 20 CFU/mL).

### Passive immunization

Female C57BL/6 mice (*n*=9-10) reared under SPF conditions (6 weeks old; Charles River Laboratories) received intraperitoneal injections of 175 µL of heat-inactivated (HI) terminal serum pooled from mice immunized with diABZI or rACP-diABZI (as described above) on days -1 and day 0 relative to infection. Mice in diestrus were implanted subcutaneously with a slow-release 17-β estradiol pellet (6.5 mg; Innovative Research of America, Cat. No. E-121) two days prior to intravaginal challenge with 10^6^ CFU of piliated Ng WHO X ^43,59^. Vaginal swabs were collected daily for 7 days and processed for CFU enumeration as described above.

### Ethics Statement

All animal work was conducted in accordance with the standards of the Association for the Assessment and Accreditation of Laboratory Animal Care. Experiments were performed under Oregon State University Institutional Animal Care and Use Committee. Animal facilities comply with the NIH Guide for the Care and Use of Laboratory Animals (Publication No. 85-23) and operate under the supervision of a full-time attending veterinarian. Mice were monitored throughout the study and euthanized at experimental endpoints by isoflurane inhalation using a regulated compressed-gas system, followed by cervical dislocation to confirm death, in accordance with institutional guidelines and the AVMA Guidelines for the Euthanasia of Animals ^61^.

### Immunoblotting

Whole-cell lysates of Ng were prepared in Laemmli SDS buffer and normalized based on OD_600_. Recombinant protein samples were prepared by protein concentration. Samples were resolved on 4%–15% Bio-Rad Criterion TGX gradient gels (Bio-Rad) and transferred onto 0.2 µM nitrocellulose membranes (Trans-Blot Turbo; Bio-Rad) using a Trans-Blot Turbo transfer system (Bio-Rad). Membranes were blocked for 1h at room temperature in 5% (w/v) non-fat dry milk in PBS containing 0.1% Tween-20 (PBST), followed by overnight incubation at 4°C with pooled terminal sera (1:5,000) or vaginal lavage samples (1:50). After washing with PBST, membranes were probed with HRP-conjugated goat anti-mouse IgG or IgA secondary antibodies (1:10,000; Southern Biotech). Blots were developed using Clarity^TM^ ECL substrate (BioRad) and visualized on a ChemiDoc^TM^ MP imaging system (BioRad).

### Enzyme-Linked Immunosorbent Assays

Endpoint binding titers were determined using a BioMek i7 liquid handler (Beckman Coulter) as previously described ^62^. Briefly, 384-well high-binding plates (Corning #3700) were coated overnight at 4°C with 2 µg/mL rACP in carbonate-biocarbonate (CBC) buffer (15 mM Na_2_CO3, 35 mM NaHCO_3_, pH 9.5; both Sigma). Plates were washed (PBS with 0.1% Tween 20; BioTek 405LS automated washer, Agilent) and blocked in CBC containing 3% non-fat dry milk (Quality Biological) and 0.1% Kathon preservative (Sigma) for 2h at room temperature. Samples were serially diluted two-fold in PBS (Invitrogen) containing 10% fetal bovine serum (FBS), 5% goat serum, 0.05% Tween-20, 1% non-fat dry milk, and 0.1% Kathon (all Sigma). Sera were assayed starting at 1:32 and vaginal wash at 1:16. Plates were incubated overnight at 4°C, washed, and incubated with HRP-conjugated detection antibodies (Southern Biotech; total IgG: catalog #1030-05, IgG1: 1070-05, IgG2c: 1078-05, IgG3: 1100-05, IgA: 1040-05) diluted 1:4,000 in PBS with 5% FBS, 5% goat serum, 0.05% Tween-20, 0.1% Kathon for 105 min at room temperature. Plates were washed, incubated with UltraTMB HRP substrate (Sigma) for 14 min, and reactions were stopped with 2N sulfuric acid. Absorbance at 450 nm was read immediately on a SpectraMAX Plus 384 (Molecular Devices), and area under the curve (AUC) was calculated in Microsoft Excel using the trapezoid rule.

### ELISpot

Female C57BL/6 mice reared under SPF conditions (4-6 weeks old, Charles River Laboratories) were vaccinated as above, without the PBS control group (in accordance with 3Rs principles). Groups (*n* = 10/group) included rACP alone, CpG+MPLA, rACP+CpG+MPLA, diABZI alone, and rACP+diABZI. Splenocytes were harvested one week after final vaccination boost, red blood cells were lysed, and splenocytes cryopreserved. Upon thawing in a 37°C water bath, splenocytes were combined with DC2.4 antigen-presenting cells at 1:10 (DC2.4:splenocyte) ratio. DC2.4 cells were pre-treated with IFNy (20 ng/mL) for 24 hours prior to assay to upregulate their immunostimulatory state and to standardize their response to antigen uptake and stimulation. Cells were plated at 250,000 splenocytes/well in in pre-wetted mouse IFNγ, IL-4, or IL-17A ELISPot plates (MabTech; Cat. #3321-4HPW-2, #3311-4HPW-2, #3521-4HPW-2, respectively) in RPMI supplemented with 10% FBS. Splenocytes were stimulated for 36 h with a custom PepMix peptide pool consisting of 23 overlapping 15-mer peptides spanning the ACP Ng FA1090 sequence lacking signal peptide (PDB: 6GQ4), synthesized by JPT Peptide Technologies GmbH. Stimulation with 50 ng/mL phorbol 12-myristate 13-acetate (PMA) and 1 ug/mL Ionomycin controlled for T-cell functionality. Peptides were designed as a contiguous scan with an 11-amino acid overlap between consecutive peptides (scan format 15/11), with a free amine N-terminus and acid C-terminus. Synthetic capping of truncated sequences was applied at each synthesis step to prevent the formation of deletion peptides, which are a primary source of false-positive T cell responses. Each peptide was purified by reverse-phase HPLC (>70% purity; C18 column, linear gradient, detection at 220 nm) and identity was confirmed by mass spectrometry. Peptides were supplied as trifluoroacetate salts, freeze-dried in amber crimp-top glass vials. For pool stimulation assays, all 23 peptides were combined into a single pool and used at 46 µg/mL total peptide or 2 µg/well (equivalent to 500 ng of each individual peptide per aliquot). Individual peptides (5 ng each) were used for epitope deconvolution assays. After 36 h of stimulation, plates were developed according to the manufacturer’s protocol. Briefly, plates were washed and incubated with biotinylated anti-mouse IFNγ, IL-4, or IL-17A detection antibody for 2 h, followed by streptavidin-HRP for 1 h after washing. Spots were visualized using BCIP/NPT-plus substrate, rinsed with water, and dried before enumeration on an AID ELISpot plate reader. To map epitope reactivity, a separate IFNγ ELISPOT was performed using each of the 23 individual ACP peptides. Immunodominant peptides were defined as those eliciting IFNγ responses exceeding the mean +2 SD of non-reactive peptides. To assess MHC-I-mediated antigen presentation, DC2.4 cells were stably transduced with a lentivector encoding ACP lacking signal peptide ^63^, co-cultured with splenocytes at a 1:10, and ELISPOT was performed as described above.

### Mapping of immunodominant ACP epitopes

Crystal structures of human C type lysozyme (PDB 7XF6) and ACP (PDB 6GQ4) were retrieved from the Protein Data Bank and curated prior to analysis. Protein–protein docking was performed using the ClusPro server with default parameters, and the top-ranked cluster was selected based on cluster size and predicted binding stability. ACP-derived peptide regions corresponding to the strongest IFN-γ ELISpot responses, as described above, were mapped to the human c type lysozyme-ACP complex model and visualized using PyMol 3.1.

### Human complement dependent serum bactericidal assay

The human complement-dependent serum bactericidal assay (hSBA) was performed as previously described ^27,38^. IgG/IgM-depleted pooled normal human serum (NHS; Pel-Freeze Biologicals; Cat. # 34010-10, Lot 34943) was used as the complement source. Heat-inactivated matched pooled human serum (HI-NHS; Cat. #34020-10, Lot 13844) served as a complement-negative control. A single lot of each serum was used across all assays to minimize complement-related variability. HI pooled or individual murine sera from immunized and control groups, as specified in the text, were serially two-fold diluted in Hanks’ Balanced Salt Solution (HBSS), starting at 1:64 to 1:32,000, for pooled sera and for individual sera, respectively. Non-piliated Ng FA1090 or WHO X were harvested from chocolate agar plates and resuspended in HBSS to ∼1×10^8^ CFU/mL. Aliquots of 1×10^3^ bacteria (40 µL) were added to wells containing serially diluted test sera. After 15 min at 37°C, 10 µL of NHS or HI-NHS was added at a final concentration of 10% (v/v) and incubated for 1 h at 37°C. Following incubation, 5 µL aliquots from each well were spotted onto chocolate agar plates and incubated overnight 37°C with 5% CO_2_ for CFU enumeration. Controls included Ng with NHS, Ng with HI-NHS, Ng with test sera and HI-NHS, and Ng alone. Percent killing was calculated by comparing CFUs recovered from Ng incubated with test sera and NHS to CFUs recovered from Ng incubated with the same sera and HI-NHS across three and two independent studies for pooled and individual sera, respectively. hSBA titers were defined as the lowest serum dilution achieving ≥50% Ng killing.

### Assessment of lysozyme-inhibitory activity

The ability of vaccine-induced antibodies to block human C-type lysozyme was assessed using pooled or individual murine sera, from mice immunized with rACP, rACP-CpG+MPLA, rACP-diABZI, or respective adjuvant-only controls (PBS, CpG+MPLA, or diABZI), as specified in the text, using the EnzChek Lysozyme Assay Kit (ThermoFisher) as previously described ^27^. Briefly, sera were incubated with human lysozyme C purified from neutrophils (HL; Sigma Aldrich) in reaction buffer (0.1M sodium phosphate, pH 7.5, 0.1M NaCl, 2mM sodium azide), followed by addition of rACP (20 µM). After 30 min at 37°C, DQ substrate was added and fluorescence was monitored kinetically (excitation 485 nm/emission 530 nm). HL alone and HL+rACP served as assay controls. Lysozyme activity was calculated from a standard curve and expressed relative to sham-immunized control sera.

### Cell lines and reagents

Unless otherwise stated, all reagents were purchased from Thermo Fisher Scientific. Human vaginal epithelial cells (VK2/E6E7; ATCC CRL-2616) and uroepithelial cells (SV-HUC-1; ATCC CRL-9520) cells were obtained from ATCC and used without additional characterization. VK2/E6E7 cells were cultured in Keratinocyte Serum-Free Medium (KSFM; Fisher Scientific) supplemented with 50 µg/mL bovine pituitary extract (BPE), 0.1 ng/mL recombinant human epidermal growth factor (EGF), and 0.4 mM calcium chloride. SV-HUC-1 cells were cultured in F-12K medium (ATCC 30-2004) supplemented with 10% heat-inactivated FBS. Human uterine fibroblasts (HUtF) were purchased (HUtF; FC-0076; Lifeline Cell Technology) were cultured in FibroLife medium (LM-001) supplemented with the Low Serum FibroLife S2 LifeFactors Kit (LS-1038) and used at passage 5. All cell types were maintained at 37°C with 5% CO_2_ and subcultured before reaching confluency.

### 3D epithelial co-culture model for evaluating antibody-mediated Ng adhesion and invasion inhibition

A 3D epithelial–fibroblast co-culture model was developed to evaluate the capacity of vaccine-induced antibodies to inhibit Ng adhesion and invasion at the mucosal surface. Heat-inactivated immune sera from mice immunized with rACP + diABZI or adjuvant-only controls were pre-incubated with WHO Ng isolates prior to infection, and the model served as a functional readout of antibody-mediated blocking activity. Human uterine fibroblasts were embedded in a GelMA hydrogel to form a supporting stromal matrix enabling epithelial barrier formation. Constructs were established on HTS Transwell-96 plates (0.11 cm² surface area, 1 µm pore size; Cat. #PSRP004R1; MilliporeSigma) and maintained until a stable epithelial barrier was established (∼7 days). To prepare the hydrogel, gelatin methacrylate (GelMA; 50% degree of substitution; Advanced BioMatrix) was dissolved per manufacturer’s instructions and a previously described protocol ^41^. Briefly, 10% (w/v) GelMA was prepared in 1×PBS, and HUtF were suspended in this solution (13,000 cells/well) to achieve a final GelMA concentration of 7% (w/v). Lithium phenyl-2,4,6-trimethylbenzoylphosphinate (LAP; Advanced BioMatrix) was used as the photoinitiator at 17 mg/mL. A 20 µL of cell–hydrogel suspension was added to the apical compartment of each transwell insert and photo-crosslinked under 405 nm light for 30 sec, per manufacturer’s recommendations. The basal compartment was filled with 235 µL of complete FibroLife medium and constructs were incubated overnight at 37°C with 5% CO₂. The following day, epithelial cells (VK2/E6E7 or SV-HUC-1) were seeded onto the hydrogel surface at densities of 30,000 cells per insert in 35 µL of a 1:1 mixture of FibroLife complete medium and the appropriate epithelial cell medium. The basal compartment was replenished with 235 µL of the same mixture. After overnight adherence at 37°C and 5% CO_2_, media were removed from both compartments. After incubation, the media were removed from both apical and basal compartments. An air–liquid interface was established by replenishing only the basal compartment (235 µL of 1:1 media mixture, replaced every 48 h) while leaving the apical compartment without media. The 3D co-culture models were maintained at 37°C with 5% CO_2_ for 7 days prior to Ng infection.

### Barrier integrity

Transepithelial electrical resistance (TEER) was measured to assess the epithelial barrier integrity prior to Ng infection ^64^. Measurements were performed using a Milicell ERS-3.0 Digital Volt-Ohmmeter (MilliporeSigma) equipped with a manual 96-well plate electrode probe (MERS0396P). TEER was recorded on days 3, 5, and 7 of epithelial differentiation. Prior to each measurement, both compartments were equilibrated to room temperature; the apical compartment were filled to 70 µL and the basal compartment to 235 µL. Background resistance was determined using inserts (no cells), and final TEER values (Ω·cm²) were calculated by subtracting background resistance and multiplying by the insert surface area (0.11 cm²). The electrode was sterilized between measurements per manufacturer instructions.

### Adhesion and invasion assays

Ng WHO X Pili^+^Opa^-^ (MOI of 10) were added to transwell inserts following 30 min pre-incubation with HI sera from mice vaccinated with diABZI or rACP+diABZI. Sera were mixed with Ng WHO X suspensions at a 1:1 ratio immediately prior to infection at final serum dilutions of 1:20, 1:40, and 1:80. Infections were carried out for 3 h at 37°C with 5% CO_2_, with 200 µL of basal media maintained throughout. Following incubation, apical were collected for downstream analyses. To quantify adhesion, cultures were washed three times with PBS to remove non-adherent Ng, lysed with 1% saponin in PBS (Sigma-Aldrich) for 15 min at room temperature, and lysates were serially diluted and plated on GC agar for CFU enumeration. To quantify Ng invasion, following the initial 3 h infection, media were replaced with epithelial cell-specific medium containing gentamicin (100 µg/mL; Sigma-Aldrich) to eliminate extracellular gonococci. After 90 min at 37°C with 5% CO_2_, cultures were washed three times with PBS and lysed with 1% saponin as described above. Lysates were serially diluted and plated on GC agar for CFU enumeration. All GC agar plates were incubated for 18-24 h at 37°C with 7% CO_2_ before colony counting.

### Integrated analysis of immune correlates of protection

Immune readouts evaluated in the 19 vaccinated animals comprised seven antibody endpoints measured by ELISA (serum IgG, IgG1, IgG2c, IgG3, and IgA; vaginal IgG and IgA), serum lysozyme rescue activity (percent recovery), and serum bactericidal antibody (hSBA) titer against the challenge strain *Neisseria gonorrhoeae* WHO X. ELISA endpoint titers were log_10_-transformed (AUC log_10_) prior to analysis to improve normality and stabilize variance. hSBA titers were determined as the reciprocal of the highest serum dilution producing ≥50% killing relative to complement-only controls. For pooled-sera analyses reported elsewhere in this manuscript, the per-pool hSBA titer is the arithmetic mean of technical replicates of a single hSBA assay. For the per-animal correlates analyses, each mouse was assayed in three independent biological replicates, and the per-mouse value used in all Cox proportional hazards, Kaplan–Meier, ROC, and group comparison analyses was the median of these three replicates. All statistical analyses were performed in R v4.5 (R Foundation for Statistical Computing). Univariate associations between continuous immune parameters and post-challenge bioburden (log_10_ CFU/mL) were assessed using Pearson correlation (linear) and Spearman rank correlation (monotonic). Coefficients are reported with two-sided p-values. The false discovery rate was controlled separately within each post-challenge timepoint using the Benjamini–Hochberg procedure. The relationship between each pre-challenge immune parameter and time to bacterial clearance was modeled by univariate Cox proportional hazards regression using the survival package. Hazard ratios are reported per one-standard-deviation increase in the standardized predictor with 95% Wald confidence intervals. The forest plot in Figure 5a was generated using the forestmodel package. Discrimination of binary Day 3 clearance by each pre-challenge immune parameter was quantified by the area under the receiver-operating characteristic curve (AUC) using the pROC package, with curves rendered using ggroc. AUC=0.5 indicates chance performance and AUC=1.0 indicates perfect discrimination. For Kaplan–Meier survival analysis, animals were stratified at a pre-challenge hSBA threshold of 2,048, the discrete two-fold dilution boundary that maximized log-rank separation between strata. Cumulative clearance curves were estimated by the Kaplan–Meier method (survival package) and plotted using survminer and ggsurvfit; the log-rank test was used to compare strata. For categorical comparisons across clearance categories, pre-challenge hSBA titers across the four ordered clearance categories (Cleared Day 3, Cleared Day 5, Cleared Day 7, Never cleared) were compared using the Kruskal-Wallis test for the overall four-group contrast and pairwise Mann-Whitney U tests for between-category comparisons. All *p*-values are two-sided; statistical significance was defined at α=0.05.

### Statistical analysis

Statistical analyses were performed using GraphPad Prism (version 11; GraphPad Software). Ng clearance rates were compared between groups using the Kaplan-Meier method with logrank (Mantel-Cox) test. Bacterial burden at each timepoint was compared using multiple Mann-Whitney tests with FDR correction (Benjamini-Hochberg method). The area under the curve (AUC) of log_10_-transformed CFU/mL over the seven-day infection period was calculated using the trapezoid rule and compared between groups by Mann-Whitney test. ELISA antibody titers (AUC) were log10-transformed prior to analysis. Data were assessed for normality using the Shapiro-Wilk test. One or more groups in each analyte violated the assumption of normality, including control groups (PBS, diABZI, adjuvant-only) where values were at the assay lower limit of detection resulting in zero variance, and non-parametric tests were therefore used throughout. Group differences were evaluated by Kruskal-Wallis test followed by Dunn’s post-hoc test with Bonferroni correction for multiple comparisons. Statistical significance is denoted as follows: \**p* < 0.05, \*\**p* < 0.01, \*\*\**p* < 0.001, \*\*\*\**p* < 0.0001. The IgG2c:IgG1 ratio was calculated for each mouse as the difference between log_10_-transformed IgG2c and IgG1 AUC values. Differences in the IgG2c:IgG1 ratio among rACP, rACP+CpG+MPLA, and rACP+diABZI groups were assessed by Kruskal-Wallis test with Dunn’s multiple comparisons post-hoc test and Bonferroni correction. Control groups (PBS, CpG+MPLA, and diABZI) were excluded from this analysis as IgG2c was undetectable in the absence of rACP antigen. hSBA titers were log₂-transformed and analyzed by two-way ANOVA with strain (FA1090 and WHO X) and treatment group as factors, followed by Tukey’s multiple comparisons test. Each treatment group was tested in three independent experiments using pooled sera. Lysozyme rescue activity was compared among treatment groups with pooled sera, as described above, by one-way ANOVA with Šídák’s multiple comparisons test. Data are expressed as mean±SEM of three independent experiments. For adhesion/invasion assays, comparisons between two groups were performed using Welch’s *t*-test when variance was unequal by F-test. Comparisons among three or more groups were performed by one-way ANOVA with Tukey’s correction or two-way ANOVA with Dunnett’s correction for multiple comparisons, as appropriate. A *p*-value of <0.05 was considered statistically significant. Sample sizes, replicates, and specific tests used are indicated in the corresponding figure legends. Peptide-specific IFNγ ELISpot responses were analyzed in GraphPad Prism by two-way ANOVA with peptide identity and adjuvant group as factors and the peptide×adjuvant interaction term included. Pairwise within-peptide comparisons of rACP+diABZI versus rACP+CpG+MPLA were corrected for multiplicity using Šídák’s method across the eight within-row contrasts (seven individual peptides plus the peptide pool).

### Visualization and illustrations

Data graphs were generated using GraphPad Prism 11 or ggplot, as described in the text. Schematic illustrations were created with BioRender.com and Adobe Illustrator.

## CONTRIBUTIONS

A.C. contributed to investigation, methodology, formal analysis, visualization, and writing—original draft. I.C. contributed to conceptualization, investigation, formal analysis, writing—original draft, and writing—review & editing. J.N. contributed to investigation, formal analysis, visualization, and writing—original draft. L.S. contributed to investigation. I.P.R. contributed to investigation, methodology, and performed experiments. J.A. contributed to investigation. C.L. contributed to formal analysis and visualization. M.A.S. contributed to investigation. A.N.M. contributed to investigation, formal analysis, and writing—original draft. V.D. contributed to conceptualization, writing—original draft, and funding acquisition. A.E.S. contributed to conceptualization, methodology, investigation, data curation, writing—original draft, writing—review & editing, formal analysis and visualization, supervision, project administration, and funding acquisition. All authors read and approved the final manuscript.

## DATA AVAILABILITY

All data supporting the findings of this study are available within the article and its Supplementary Information files. The source data underlying Figs. 1–5 and Supplementary Figs. 1–16 are provided as a Source Data file.

## CODE AVAILABILITY

Custom R scripts used for survival analysis, ROC analysis, and correlates of protection are provided in DATA SOURCE.

## ACKNOWLEDGEMENTS

We thank Lixin Li for technical assistance with animal experiments. Funding was provided to Dr. Aleksandra Sikora through grant R01-AI117235 and to Dr. DeFilippis through grant R01-AI177293 from the National Institute of Allergy and Infectious Diseases, National Institutes of Health. ELISAs were conducted in the Regional Biocontainment Laboratory at Duke University, which received partial support for construction and renovation from the National Institutes of Health (UC6-AI058607 and G20-AI167200), and facility support from the National Institutes of Health (UC7-AI180254). The funder had no role in study design, data collection or analysis, publication, or manuscript preparation.

## SUPPLEMENTARY INFORMATION

Supplementary materials are included as a PDF file.

