## Supplemental Material for "Mucosal ACP subunit vaccine formulated with a small molecule STING agonist drives protective T_H_1/T_H_17 responses and serum bactericidal antibodies against *Neisseria gonorrhoeae*"

**Supplementary information**

**Supplementary Table S1:**

| Variable | HR | 95% CI | p-value |
|---|---|---|---|
| hSBA titre | 1.00 | 1.00 – 1.00 | <0.001*** |
| Serum IgA | 1.91 | 1.09 – 3.37 | 0.025* |
| Serum IgG | 131.12 | 0.27 – 63,454.90 | 0.122 |
| Serum IgG3 | 1.58 | 0.77 – 3.23 | 0.213 |
| Serum IgG2c | 1.54 | 0.41 – 5.74 | 0.519 |
| Vaginal IgA | 0.94 | 0.66 – 1.32 | 0.711 |
| Lysozyme (%) | 1.00 | 0.99 – 1.02 | 0.815 |
| Serum IgG1 | 1.97 | 0.00 – 1,333.03 | 0.838 |
| Vaginal IgG | 1.04 | 0.64 – 1.69 | 0.887 |

**Supplementary Table S1:** Inline table showing values of Hazard Ratio (HR) per one standard deviation increase with 95% confidence intervals (95% CI) and *p*-value for each parameter in support for Fig 5a.

### Supplementary Fig. S1

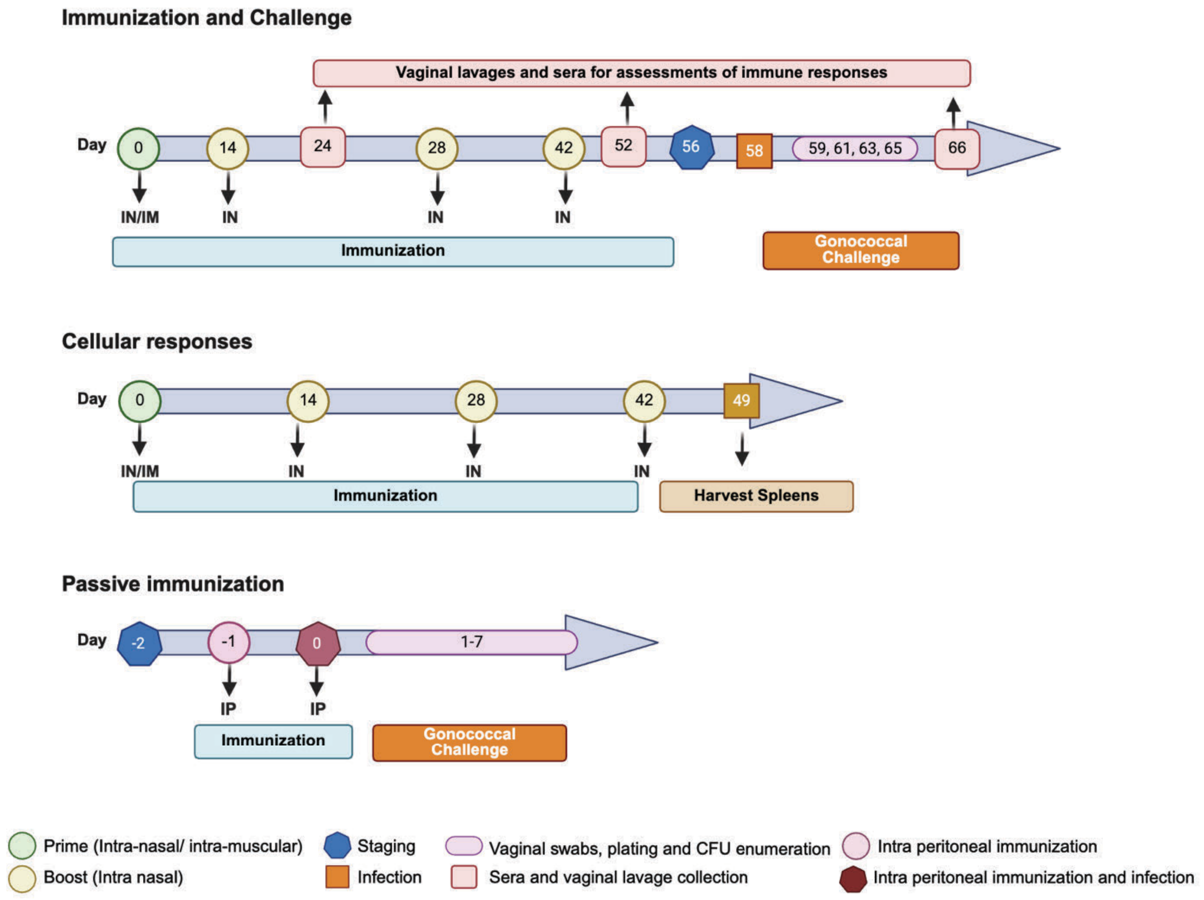


**Supplementary Figure S1. Experimental timelines for immunization, challenge and evaluation of humoral and cellular correlates of protection.**

a) **Immunization and challenge.** In the primary study, female C57BL/6 mice (*n*=17-22/group) received one prime followed by three booster doses of immunization at an interval of two weeks. Sera and vaginal lavages were collected 10 days after each boost for assessment of immune responses. Mice were staged two weeks after the last booster dose and challenged intravaginally two days later (Day 58) with *N. gonorrhea* to track the bacterial bioburden and clearance kinetics.

b) **Cellular responses.** A parallel correlates study (*n*=10 mice/group) employed a same immunization schedule, where spleens were harvested a week after the final booster for cellular analysis.

c) **Passive immunization.** Recipient mice (*n*=9-10/group) were intraperitonially administered pooled sera of diABZI and ACP-diABZI groups from the previous experiments on days −2 and −1 prior to gonococcal challenge to determine the contribution of humoral immunity. Following challenge, infection progression was monitored over a 7-day period.

Graphical elements denote timing of immunization, sampling, challenge, and downstream analyses.

**Supplementary Fig. S2**

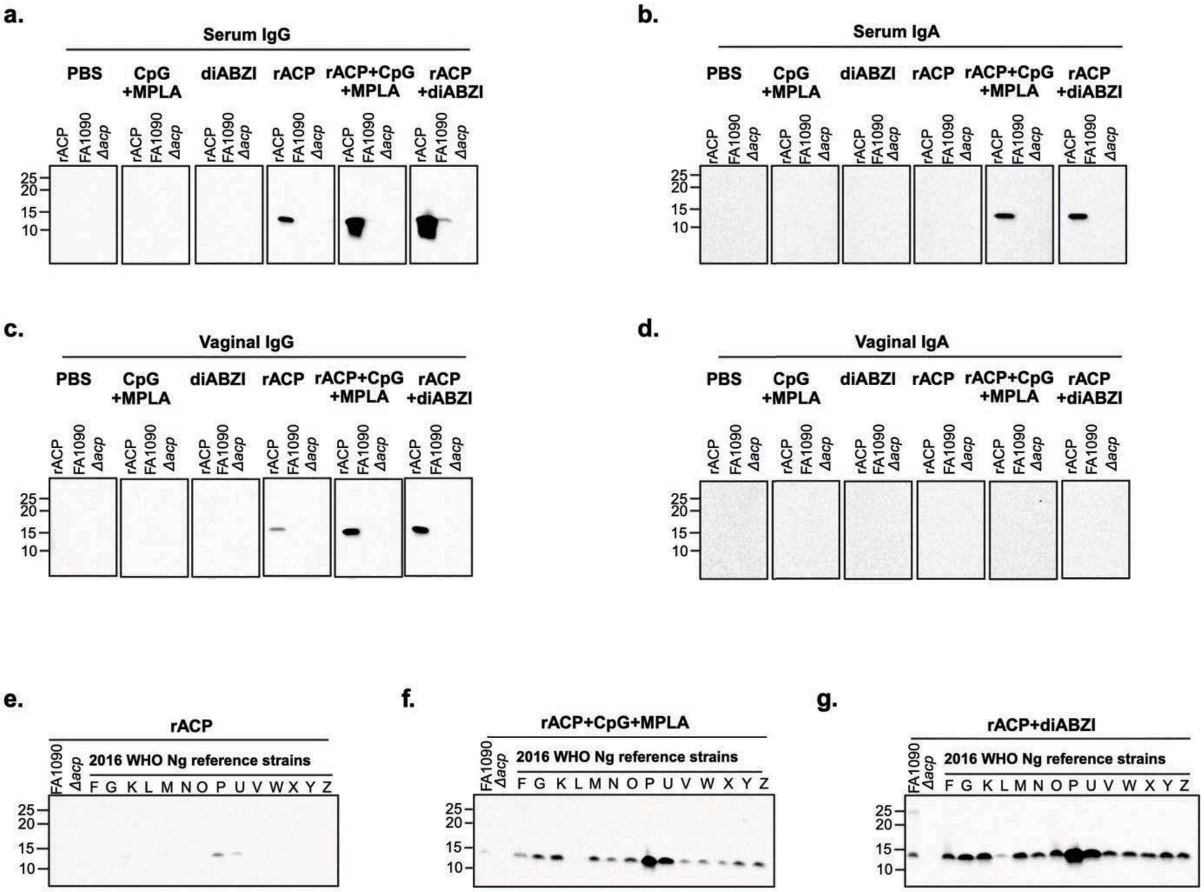


**Supplementary Figure S2. Immunoblot analysis of systemic and mucosal antibody responses induced by ACP-based vaccines.** Pooled sera and vaginal lavages obtained from the terminal samples were used as primary antibodies to probe purified recombinant ACP (rACP) and gonococcal whole-cell lysates. **Panels a,b** show antigen-specific systemic IgG and IgA reactivity, respectively and **Panel c,d** demonstrate mucosal IgG and IgA responses respectively. **Panel e-g** illustrate total serum IgG reactivity against whole-cell lysates from diverse whole-cell lysates from 14 genetically and temporally diverse Ng strains in the 2016 WHO Ng reference panel, demonstrating cross-strain antigen recognition using sera from rACP, ACP-CpG+MPLA and ACP-diABZI immunized mice. All the blots for each immunoglobulin isotype were run in the same gel, processed and imaged using identical acquisition parameters.

**Supplementary Fig. S3**

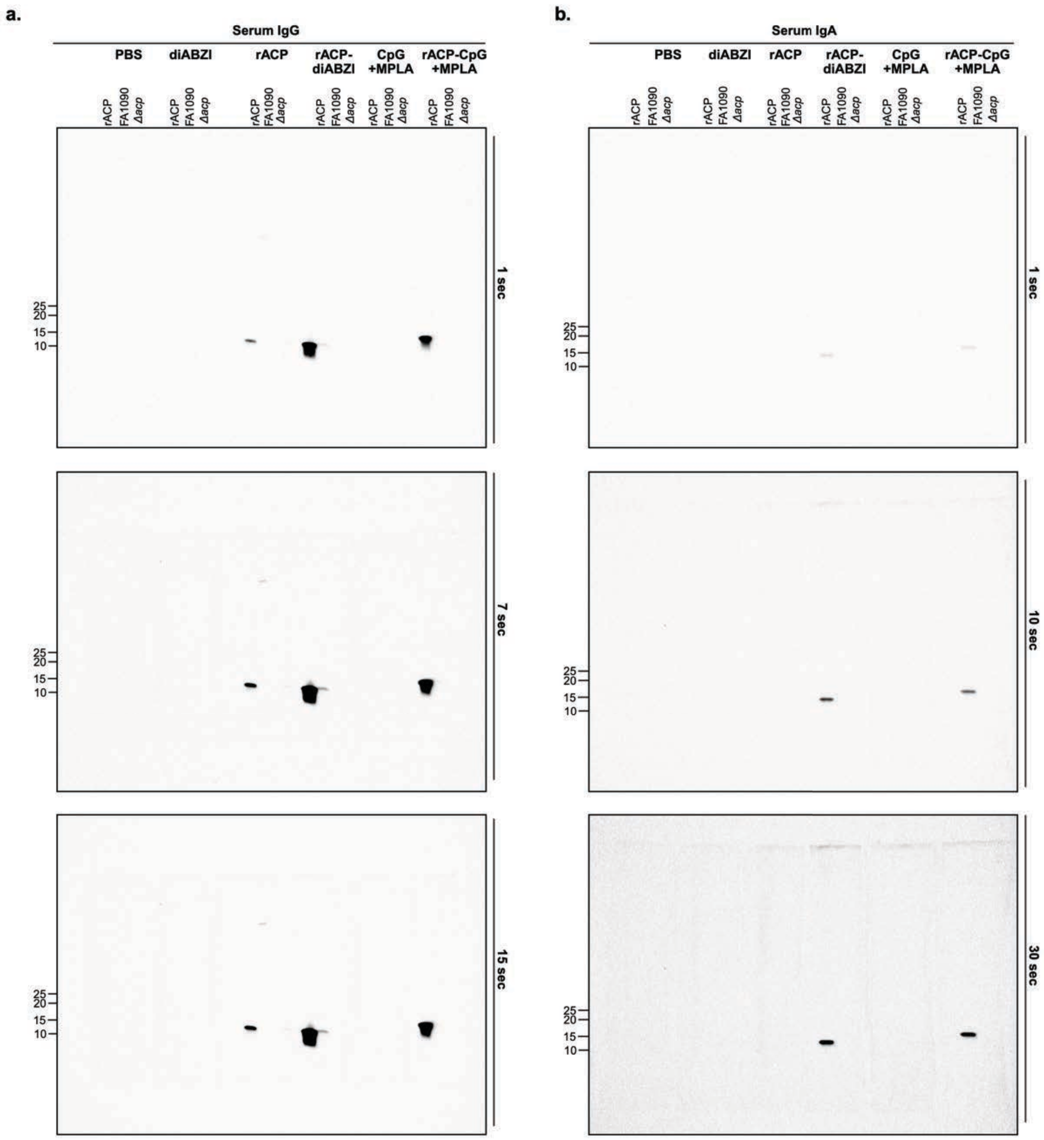


**Supplementary Figure S3**. **Uncropped and unprocessed sera immunoblot images supporting Supplementary Figure S2 a & b.** Original immunoblots at three different exposure times indicated on the right**.** Pooled sera from the terminally bled female BL57/c mice immunized with recombinant ACP (rACP) or adjuvanted with CpG+MPLA (rACP+CpG+MPLA) or diABZI (rACP-diABZI) alongside respective controls; served as primary antibody in immunoblotting assays to detect ACP specific serum **(a)** IgG and **(b)** IgA. Molecular weight (in kDa) is indicated to the left of each panel.

**Supplementary Fig. S4**

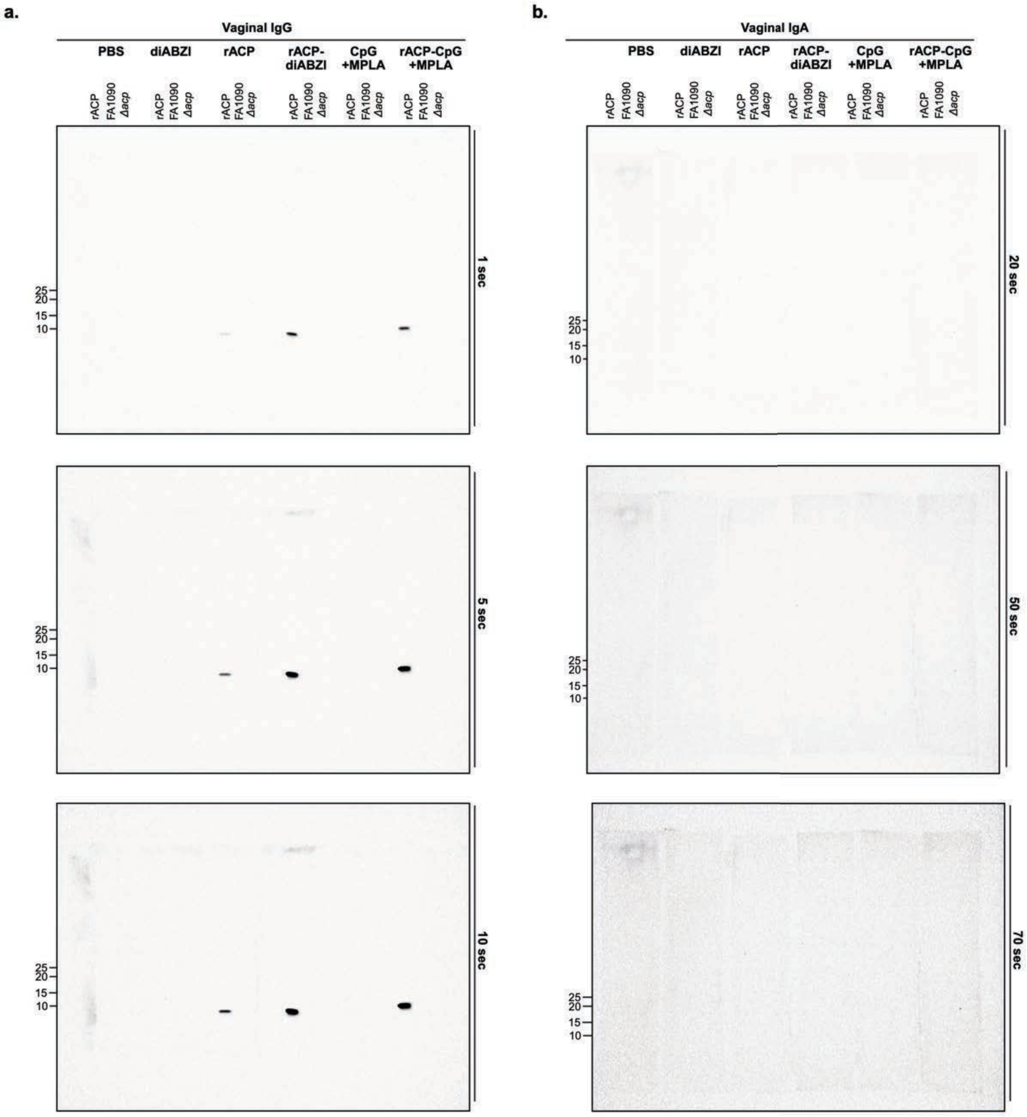


**Supplementary Figure S4. Unprocessed and uncropped vaginal wash immunoblots supporting Supplementary Figure S2 c & d.** Pannels present uncropped and original immunoblots corresponding to **Supplementary Fig. S2c** (Vaginal IgG) and **Supplementary Fig. S2d** (Vaginal IgA) at three different exposure times indicated on the right side of the figure**.** Assay utilized pooled terminal vaginal lavages from female BL57/c mice immunized with recombinant ACP (rACP), rACP combined with CpG+MPLA (rACP+CpG+MPLA), rACP combined with diABZI (rACP-diABZI) and CpG+MPLA, diABZI or PBS as a control. The molecular weight of protein markers (in kDa) is indicated to the left of each panel.

**Supplementary Fig. S5**

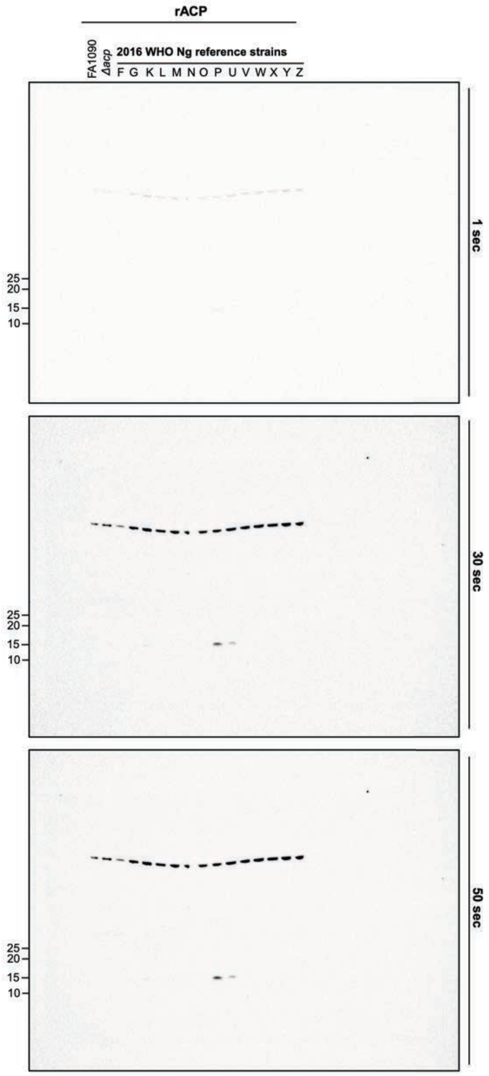

### Supplementary Fig. S6

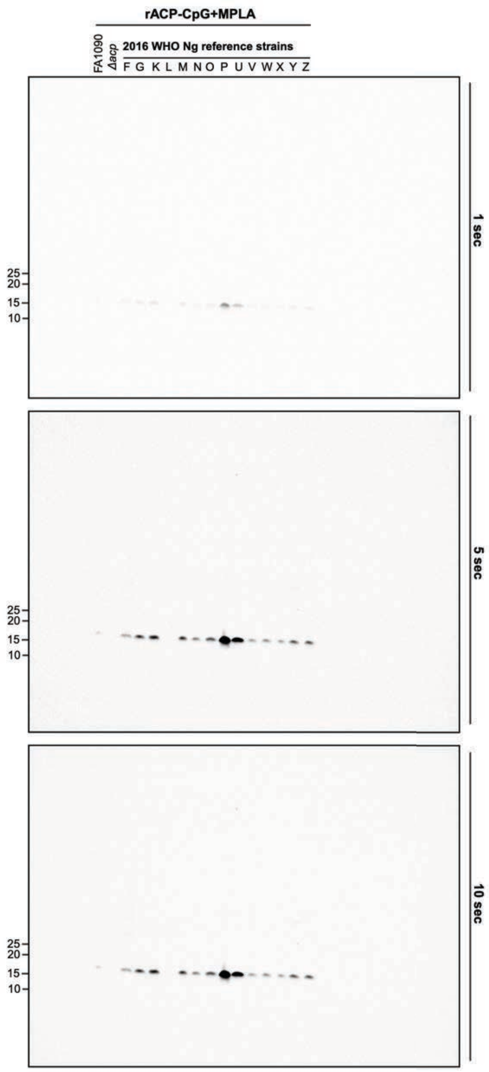

**Supplementary Fig. S7**

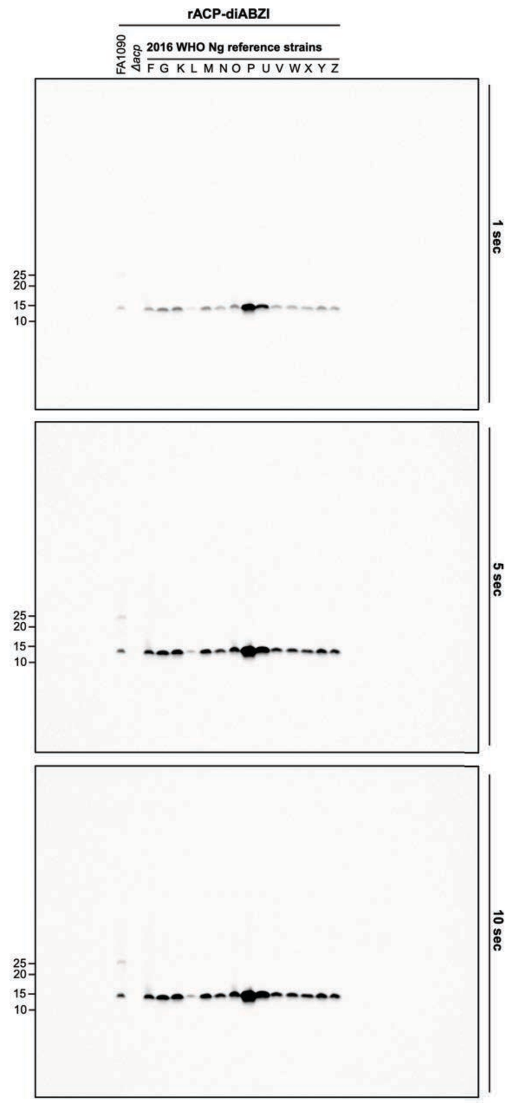


**Supplementary Figures S5-7. Original, uncropped, and unprocessed immunoblot images displaying serum IgG cross-reactivity to diverse Ng whole cell lysate supporting Supplementary Figure S2 e-g.** Three replicate uncropped immunoblot images corresponding to the exposure conditions indicated on right. Pooled terminal sera from female BL57/c mice vaccinated with rACP alone **(Supplementary Fig. S5)** or adjuvated with CpG+MPLA **(Supplementary Fig. S6)** or diABZI **(Supplementary Fig. S7)**; were used to evaluate serum IgG binding to whole cell lysate from *Neisseria gonorrhoeae* (Ng) FA1090, isogenic *Δacp* mutant, and the genetically and temporally diverse 2016 WHO Ng reference strains (F, G, K, L, M, N, O, P, U, V, W, X, Y, Z). Molecular weight (in kDa) is indicated to the left of each panel.

### Supplementary Fig. S8

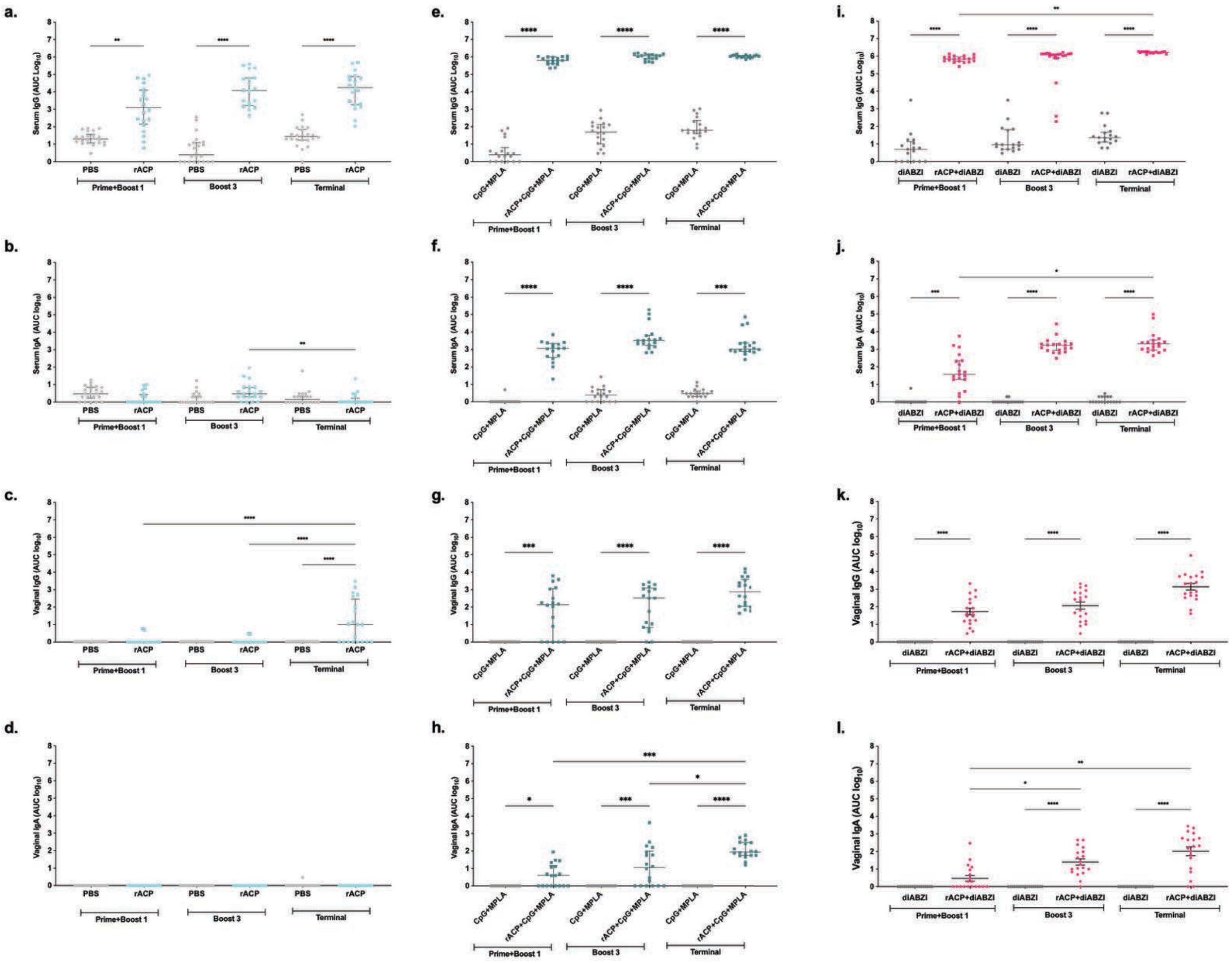


**Supplementary Fig. S8: Longitudinal systemic and mucosal antigen-specific IgG and IgA responses induced by ACP vaccination across three adjuvant formulations.** Mice were immunized intranasally with ACP formulated with PBS,

CpG/MPLA, or diABZI, and serum and vaginal lavage samples were collected 10 days after the second and fourth immunization as well as at the termination of the study. The samples were analyzed by ELISA for ACP-specific IgG and IgA response. Serum IgG and IgA **(a, b)** and vaginal IgG and IgA **(c, d)** in PBS- versus ACP-immunized mice. Serum IgG and IgA **(e, f)** and vaginal IgG and IgA **(g, h)** in mice receiving CpG/MPLA adjuvant alone versus ACP+CpG/MPLA. Serum IgG and IgA **(i, j)** and vaginal IgG and IgA **(k, l)** in mice receiving diABZI alone versus ACP+diABZI. Each symbol represents one animal; horizontal bars indicate group medians with interquartile range. Statistical comparisons by Kruskal-Wallis test with Dunn's post-hoc correction: *$p<0.05$, **$p<0.01$, ***$p<0.001$, ****$p<0.0001$.

**Supplementary Fig. S9**

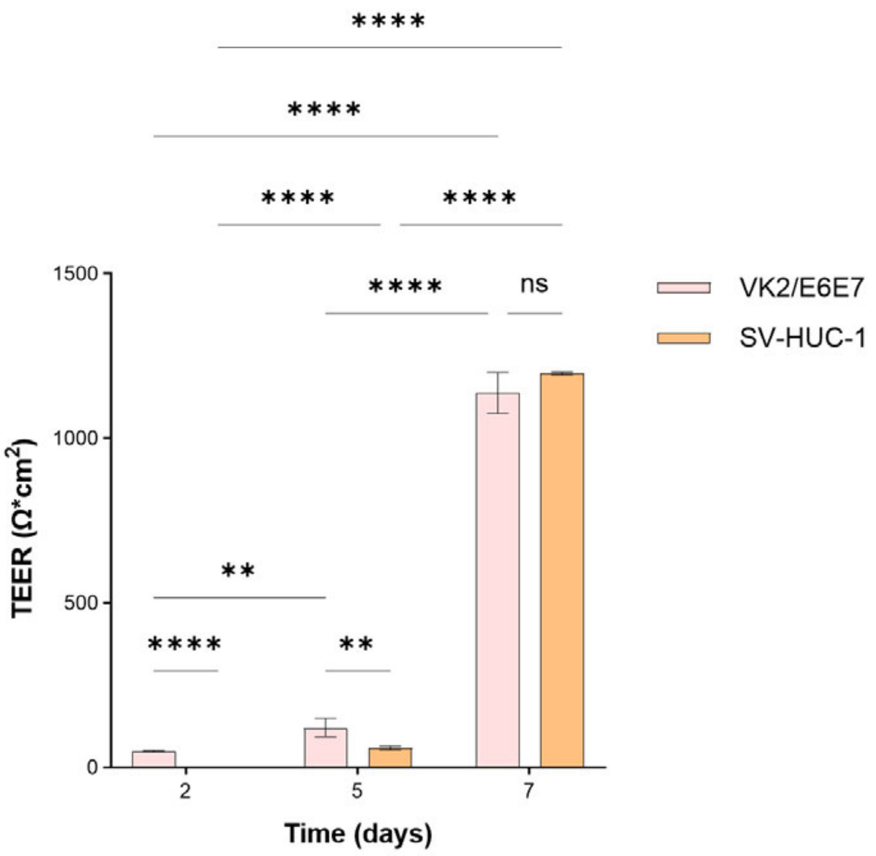


**Supplementary Figure S9.** Comparison of barrier integrity in 3D mucosal models. Transepithelial electrical resistance (TEER) was measured in 3D models prior to infection with *N. gonorrhoeae* strain WHO X. Models comprised vaginal epithelial cells (VK2/E6E7, female) or urothelial cells (SV-HUC-1, male) seeded on top of human fibroblasts encapsulated in GelMA (10% w/v). $n$=6; **$p$<0.01, ***$p$<0.001, ****$p$<0.0001; Two-way ANOVA with Tukey's post hoc test.

**Supplementary Fig. S10**

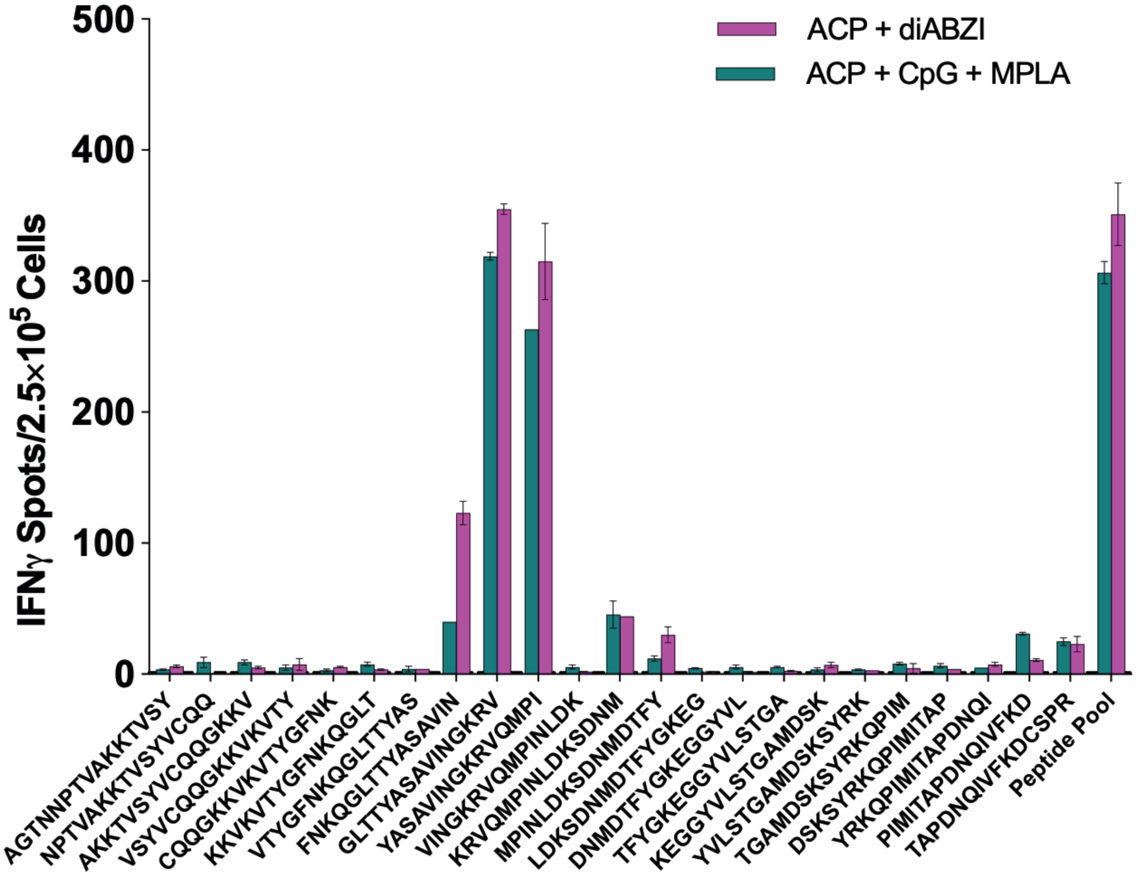


**Supplementary Figure S10. ACP Epitope Deconvolution by IFN-γ ELISpot.** Splenocytes harvested one week after final vaccination boost from female C57BL/6 mice (*n*=10/group) vaccinated with rACP+diABZI or rACP+CpG+MPLA were stimulated with each of 23 individual overlapping 15-mer ACP peptides spanning the *Neisseria gonorrhoeae* FA1090 sequence (scan format 15/11 with 11-amino acid overlap) to map epitope-specific IFN-γ responses. Splenocytes were co-cultured with DC2.4 antigen-presenting cells (1:10 ratio) and plated at 250,000 cells/well in IFN-γ ELISpot plates stimulated for 18 hours with individual peptides (0.5 mg each). Spots were enumerated following plate development and are expressed as IFN-γ-secreting cells per $2.5 \times 10^5$ splenocytes. Immunodominant peptides were defined as those eliciting IFN-γ responses exceeding the mean +2 standard deviations of non-reactive peptides. Both vaccine formulations elicited robust IFN-γ responses against peptides within the N-terminal and C-terminal regions of ACP, with the rACP+diABZI formulation demonstrating notably higher responses to immunodominant epitopes compared to the CpG+MPLA adjuvant combination. Error bars represent standard error of the mean. Data are representative of two independent experiments.

### Supplementary Fig. S11

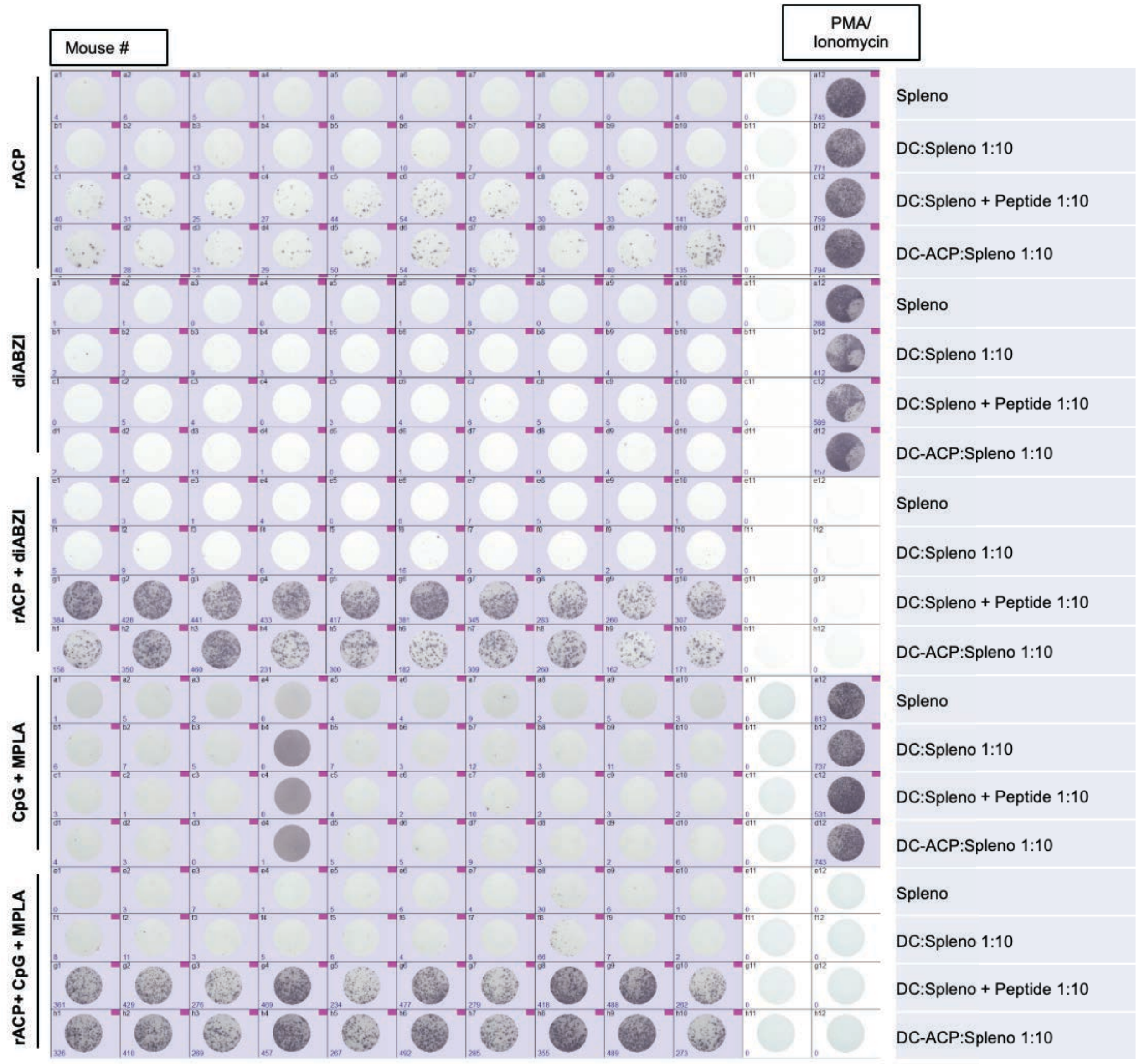


**Supplementary Fig. S11. Representative raw ELISpot plate images showing antigen-specific cellular responses across immunization groups supporting Figure 3a and 3b**. ELISpot assays were performed using splenocytes isolated from female C57BL/6 mice (n=10/group) immunized with only adjuvants (CpG+MPLA and diABZI) or vaccines (rACP, rACP+CpG+MPLA,, and rACP+diABZI). Splenocytes were harvested one week following the final immunization boost, cryopreserved, thawed, and co-cultured with DC2.4 antigen-presenting cells at a 1:10 ratio (DC2.4: splenocytes). Cells were plated at 2.5 × $10^5$ splenocytes per well in pre-coated INFg ELISpot plates.

Each row corresponds to an individual mouse, and columns represent replicate wells under the indicated stimulation conditions: unstimulated splenocytes (Spleno), DC2.4 co-culture (DC:Spleno), ACP peptide pool-stimulated co-culture (DC:Spleno + peptide) (**Fig. 3a)**, and DC2.4 cells expressing ACP (DC-ACP:Spleno) **(Fig. 3b)**. Positive control wells stimulated with PMA/ionomycin are shown in the rightmost column. Spot-forming units (SFUs) were visualized using BCIP/NPT substrate and enumerated using an automated ELISpot reader. Images shown are unprocessed representative plate scans. Minor cropping was applied uniformly to align wells; no selective enhancement was performed.

**Supplementary Fig. S12**

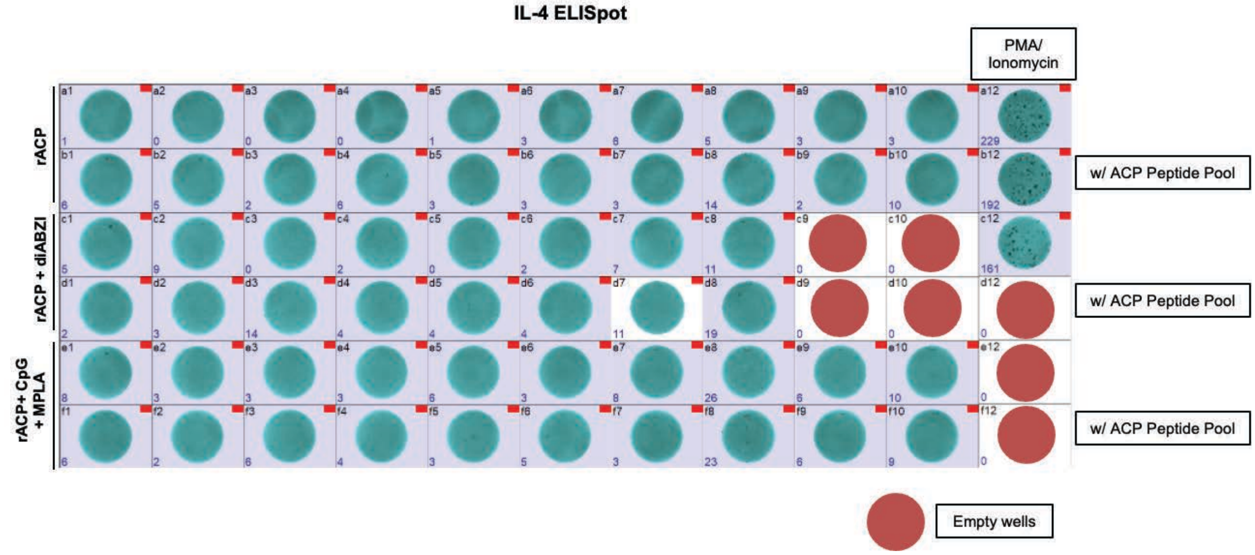


**Supplementary Fig. S12. Raw IL-4 ELISpot plate images showing Th2 cytokine responses in support to Fig. 3c.** Splenocytes isolated from immunized mice (only vaccine groups) were co-cultured with DC2.4 antigen-presenting cells at a 1:10 ratio and stimulated for 18 h with an overlapping ACP peptide pool or left unstimulated. Cells were plated at $2.5 \times 10^5$ splenocytes per well in IL-4-specific ELISpot plates. Each well corresponds to an individual mouse replicate under defined stimulation conditions for the designated vaccine groups on the left. PMA/ionomycin-treated wells served as positive controls. Images shown are representative of raw plate scans used for quantification.

**Supplementary Fig. S13**

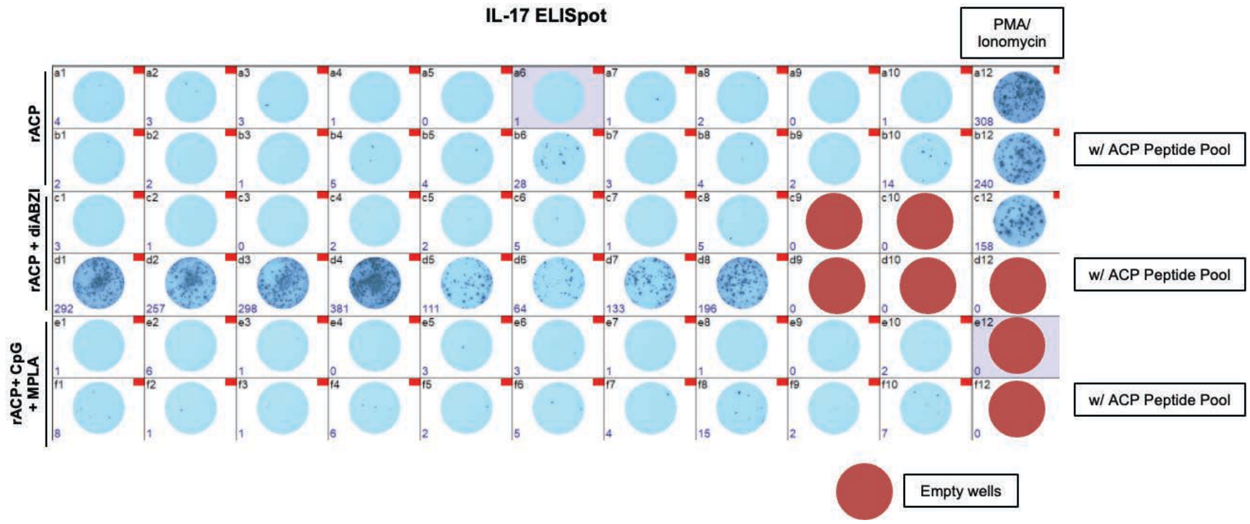


**Supplementary Fig. S13. Raw IL-17A ELISpot plate images showing Th17 cytokine responses supporting Figure 3d.** IL-17A secretion was measured in splenocytes from immunized mice following co-culture with DC2.4 antigen-presenting cells and stimulation with an ACP-derived peptide pool. Each well contains $2.5\times10^5$ splenocytes, and each well corresponds to an individual animal within the indicated immunization group (on left). Spot counts (displayed numerically) represent IL-17A–producing cells after 18 h stimulation. PMA/ionomycin-treated wells serve as positive controls.

**Supplementary Fig. S14**

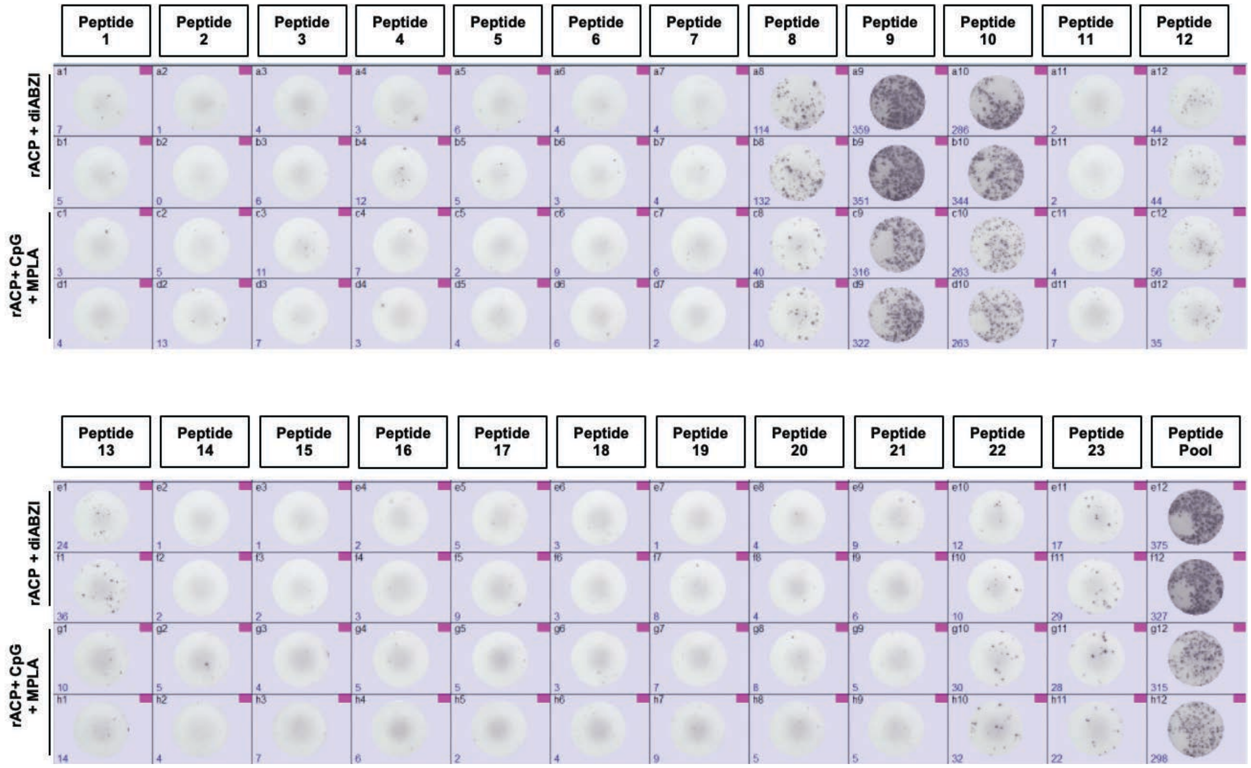


**Supplementary Fig. S14. Raw ELISpot images in support of Fig. 3e for the identification of antigen-specific T-cell epitopes across ACP peptide library.** To define immunodominant T cell epitopes, splenocytes from immunized mice were stimulated with individual ACP-derived peptides or pooled peptide mixture. Cells were co-cultured with DC2.4 antigen-presenting cells and plated at $2.5 \times 10^5$ splenocytes per well. Each column corresponds to a distinct peptide, and each row represents an individual mouse within the indicated vaccine group (designated on the left). Spots represent IFN-γ–secreting cells following 18 h stimulation. Images shown are raw ELISpot scans prior to quantitative analysis.

**Supplementary Fig. S15**

**a**

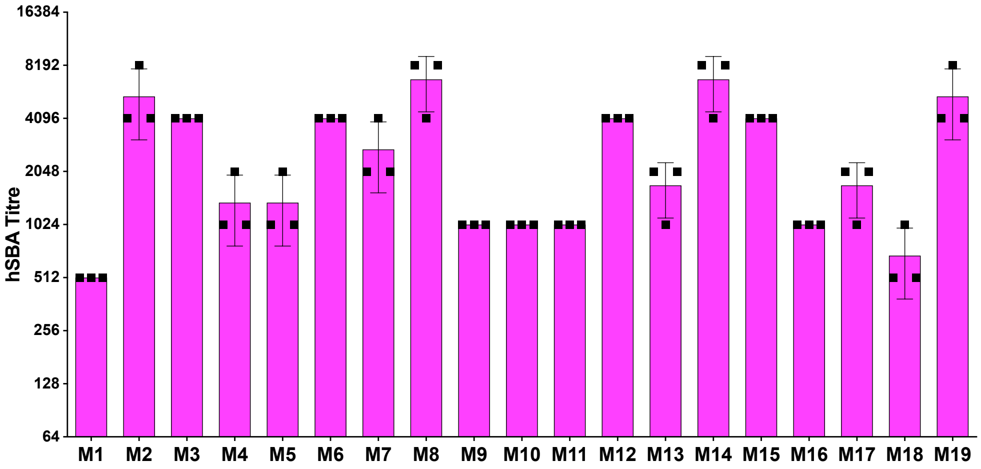


**b.**

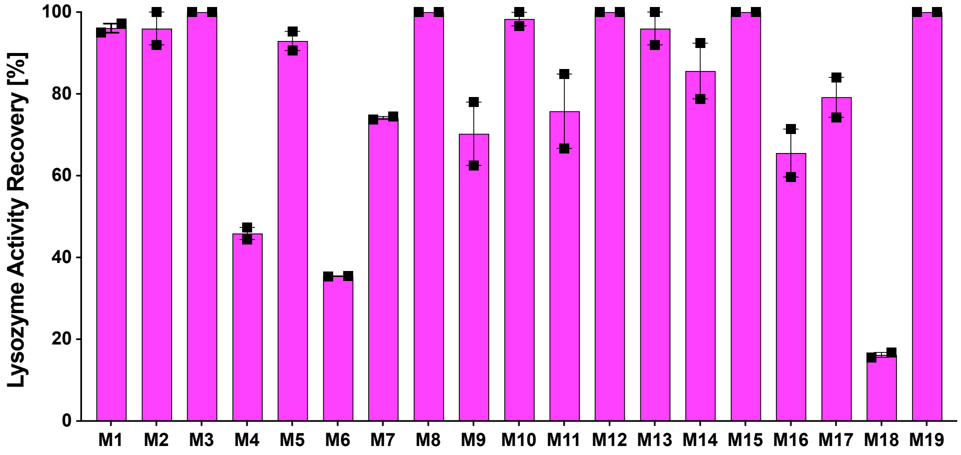


**Supplementary Figure S15. Variability of functional antibody responses within individual mice immunized with rACP-diABZI vaccine. (a)** Human serum bactericidal activity (hSBA) titers measured in terminal sera from individual mice immunized with rACP+diABZI. Each bar represents individual animal, and the point represents number of replicates performed (*n*=3/mice). Bars show mean±SEM. Titers reflect the highest serum dilution achieving ≥50% killing of *N. gonorrhoeae* WHO X. **(b)** Rescue of human lysozyme C (HL) enzymatic activity by vaccine-elicited antibodies. Terminal sera from individual mice immunized by rACP+diABZI were incubated with HL in black 96-well plates, followed by addition of rACP to inhibit HL. After 30 min, hydrolytic activity was initiated using DQ-lysozyme substrate, and fluorescence was measured kinetically at 37°C (Ex 485 nm/Em 528 nm). HL activity was quantified using a standard curve, and % rescue was calculated relative to positive (HL alone) and negative (HL + rACP) controls. Each bar represents one animal; each point shows individual replicates performed (*n*=2/mice) and bars show mean±SEM.

### Supplementary Figure S16.

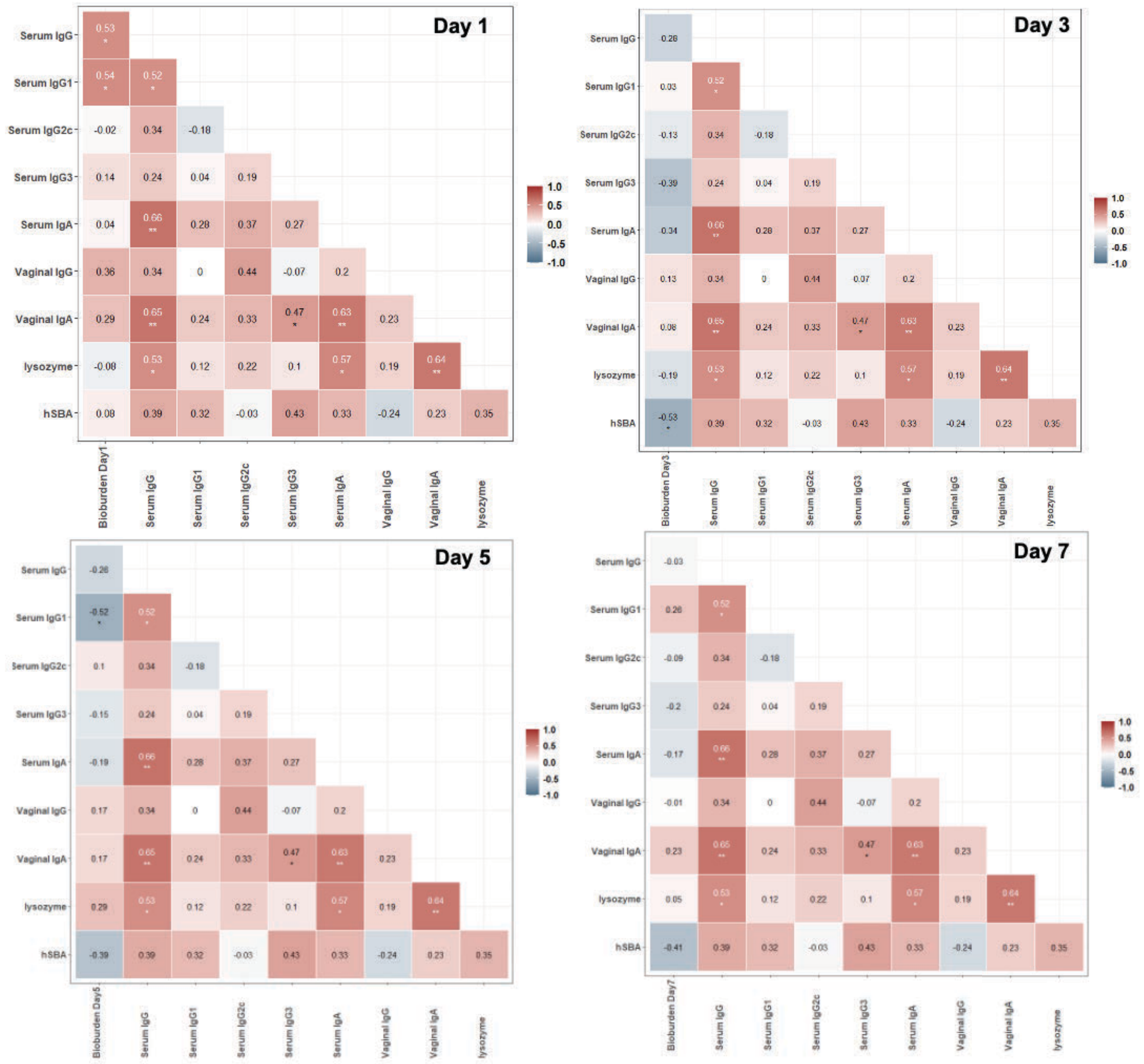


**Supplementary Figure S16.** Spearman correlation heatmaps for Days 1, 3, 5, and 7 showing all immune parameters and bioburden at each timepoint, with significance asterisks.
